# A nascent polypeptide sequence modulates DnaA translation elongation in response to nutrient availability

**DOI:** 10.1101/2021.03.31.437742

**Authors:** Michele Felletti, Cedric Romilly, E. Gerhart H. Wagner, Kristina Jonas

## Abstract

The ability to regulate DNA replication initiation in response to changing nutrient conditions is an important feature of most cell types. In bacteria, DNA replication is triggered by the initiator protein DnaA, which has long been suggested to respond to nutritional changes, nevertheless the underlying mechanisms remain poorly understood. Here, we report a novel mechanism that adjusts DnaA synthesis in response to nutrient availability in *Caulobacter crescentus*. By performing a detailed biochemical and genetic analysis of the *dnaA* mRNA we identified a sequence downstream of the *dnaA* start codon that inhibits DnaA translation elongation upon carbon exhaustion. Our data show that the corresponding peptide sequence, but not the mRNA secondary structure or the codon choice, are critical for this response, suggesting that specific amino acids in the growing DnaA nascent chain tune translational efficiency. Our study provides new insights into DnaA regulation and highlights the importance of translation elongation as a regulatory target. We propose that nascent chain sequences, like the one described, might constitute a general strategy for modulating the synthesis rate of specific proteins under changing conditions.

## INTRODUCTION

Most cells must be able to integrate external information with proliferative functions such as DNA replication, cell division, and protein synthesis. In particular, unicellular bacteria that frequently face drastic environmental changes in their natural habitat are able to precisely adjust their growth and division rate in response to environmental cues. While most bacteria proliferate rapidly under optimal conditions, they slow down growth and often enter a non-growing state under adverse conditions, for example nutrient limitation^1^. This dynamic switching between proliferative and non-growing modes contributes to stress and antibiotic tolerance, as well as the virulence of pathogenic bacteria^2–4^. Previous research has focussed on the mechanisms that allow bacteria to globally reprogram gene transcription in response to nutrient availability^5, 6^. More recently, the advance of ribosome profiling technologies has also revealed a central role of translational regulation in the adaptation to changing nutrient conditions^7–10^. Despite significant progress, the precise molecular mechanisms regulating the translation of specific proteins during nutrient limitation are incompletely understood.

When entering growth arrest, most bacteria inhibit DNA replication initiation and halt their cell division cycle with a reduced number of fully replicated chromosomes^11^. Shutting down DNA replication prior to the cessation of key metabolic functions and cell division likely preserves genome and cellular integrity. In nearly all bacteria, replication initiation depends on DnaA, an ATP-binding protein consisting of four distinct structural and functional domains: (I) a helicase loading domain, (II) a linker domain, (III) an AAA+ (ATPase Associated with diverse cellular Activities) domain, and (IV) a DNA-binding domain^12^. DnaA is only active in the ATP-bound state and, upon binding to specific DnaA-boxes in the bacterial origin of replication, it unwinds double stranded DNA and recruits the replisome. ATP hydrolysis following replication initiation subsequently inactivates DnaA^13, 14^. Due to its critical function in replication initiation, DnaA has been proposed to be a crucial player in the nutritional control of DNA replication for more than 50 years^15^. Yet, the precise molecular mechanisms that regulate this important protein in response to changing nutrient conditions remain poorly understood.

In the freshwater bacterium *Caulobacter crescentus* (*Caulobacter* hereafter), DnaA is cleared at the onset of carbon starvation when growth rate declines, which entails a block of DNA replication initiation^16–18^. *Caulobacter* is an important model for bacterial cell cycle studies due to its asymmetric cell cycle and its genetical tractability^19^. Furthermore, as a member of the *Alphaproteobacteria*, it is representative of a diverse group of bacteria, including important pathogens and agriculturally relevant species^19^. In its natural habitat, *Caulobacter* frequently experiences fluctuations in nutrient availability, for example as an indirect consequence of seasonality or changes in the microbial composition of its niche^20^.

*Caulobacter* DnaA is a relatively unstable protein that is degraded by the AAA+ protease Lon^21–23^. Although Lon-dependent proteolysis is required for rapid clearance of DnaA at the onset of starvation, the rate of proteolysis is not significantly affected by changing nutrient conditions^18^. Instead, a starvation-induced decrease in DnaA translation rate was shown to be responsible for nutritional regulation of DnaA levels^18^. However, the underlying molecular mechanism remains elusive.

Here, we reveal the mechanistic basis underlying the starvation-induced downregulation of DnaA translation in *Caulobacter*. An in-depth biochemical and genetic characterisation of the *dnaA* mRNA showed that its 5’ untranslated leader region (5’UTR) is dispensable for this regulation. Instead, inhibition of DnaA translation occurs during polypeptide chain elongation in a starvation-dependent manner, within a coding sequence downstream of the start codon. Our data are consistent with a model in which specific amino acids in the N-terminus of DnaA mediate a translation arrest in response to nutrient limitation while, or after, being added to the growing nascent polypeptide chain. In addition to providing new molecular insights into the nutritional regulation of DnaA, our findings illustrate how specific sequence motifs in nascent polypeptides can tune the first stages of translation elongation under changing nutrient conditions.

## RESULTS

### The 5’UTR of the *Caulobacter dnaA* mRNA adopts a complex secondary structure

In *Caulobacter*, DnaA levels decrease when cells are shifted from a glucose-supplemented minimal medium to the glucose-limiting medium M2G_1/10_ (Fig. 1A)^18^. Previous work established that this carbon starvation-induced downregulation of DnaA abundance is caused by decreased *dnaA* translation in combination with constitutive DnaA degradation by Lon^18^. To elucidate the mechanistic basis for starvation-dependent downregulation of DnaA, we structurally and functionally characterised the mRNA region comprising the 155 nucleotides (nt) long 5’UTR and the first 118 nt of the *dnaA* coding region (total RNA length 273 nt). Native polyacrylamide gel electrophoretic analysis of this *in vitro* transcribed ^32^P-5’-end-labelled RNA indicated a dominant conformation of the *dnaA* mRNA leader (Supplementary Fig. 1A). To determine its secondary structure, we combined mFold and ViennaRNA computational predictions with structural probing (Supplementary Fig. 1B-D). The latter was performed with Pb^2+^ to induce RNA cleavages in single-stranded regions, RNase T1 to cleave after single-stranded G residues, and RNase V1 for mapping double-stranded stretches. The experimental and *in silico* analyses suggested seven base-paired segments: one helical element (P1) and six stem-loop motifs (P2-P7, Fig. 1B). The 5’-most 8 nt are part of helix P1, which is connected via a three-way junction to stem-loops P2 and P3. The stable GC-rich stem P4 carries a G-rich loop which is highly susceptible to T1 RNase cleavage. A fifth stem-loop structure (P5) is located upstream of the AUG start codon. Two mismatches in this stem confer some instability (ΔG_folding_ of −7 kcal/mol). The region around the start codon appears mostly unstructured, suggested by T1 cleavages at nucleotides G154, G158, and G164 (Fig. 1C). The mRNA region encompassing the first 118 nt of the *dnaA* coding region folds into two stable stem-loops, P6 and P7. The additional RNase V1 cleavages in this portion of the RNA might indicate base-pairing of the 3’ terminal nucleotides with the mRNA region around the start codon.

**Fig. 1.**
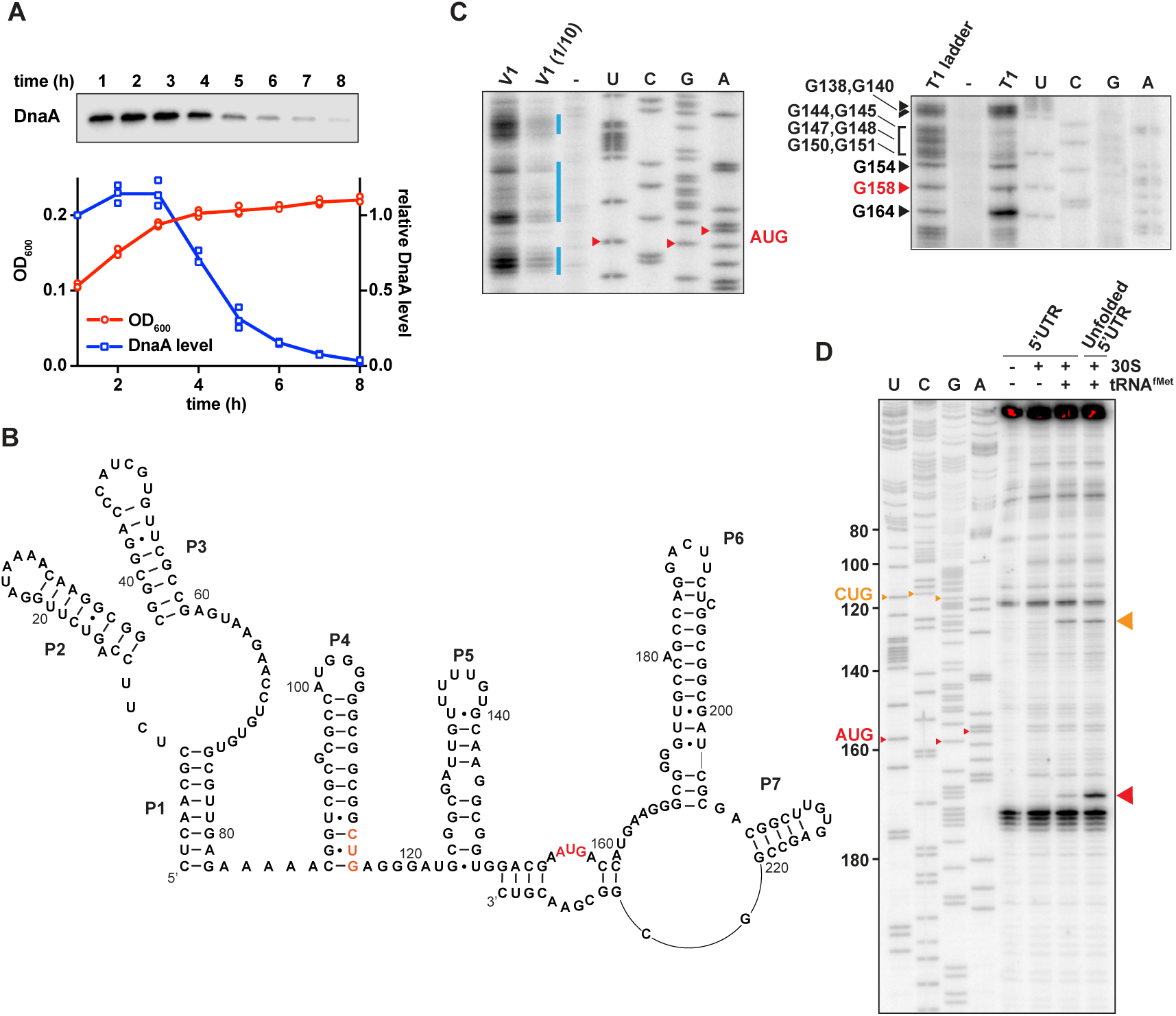
*Caulobacter dnaA* mRNA assumes a complex secondary structure that inhibits pre-initiation complex formation. (**A**) Growth curve (red) and DnaA abundance (blue) measured during a carbon exhaustion experiment. Wild type *Caulobacter* was grown in a defined minimal medium (M2G_1/10_) containing 0.02% glucose as the sole carbon source. DnaA levels were determined by Western blot and band intensities quantified relative to the first time point (t = 1 h). Averages of three independent replicates are shown with error bars representing standard errors. (**B**) Secondary structure of the mRNA region encompassing the 5’UTR and the first 26 codons of the *dnaA* open reading frame. The reported AUG start codon and the alternative CUG codon are highlighted in red and orange respectively. See Supplementary Fig. 2A for an alternative secondary structure differing in the way the 3’-terminal RNA portion folds back on the start codon region. (**C**) RNA probing gels showing the *dnaA* mRNA region containing the translation start site and the RBS. Upon partially digesting a synthetic *in vitro* transcribed RNA with V1 (left gel) or T1 (right gel) RNases, primer extension (^32^P-5’-end-labelled primer B – Supplementary Fig. 2A) was used to detect cleavage positions. V1 and V1 (1/10) lanes, samples treated with different concentrations of RNase V1. T1 lane, sample treated with T1 RNase. T1 ladder lane, RNase T1 treatment under denaturing conditions. -, mock-treated RNA. U, C, G, A lanes, sequencing reactions using the ^32^P-5’-end-labelled primer B. The bands corresponding to the start codon nucleotides are indicated in red. The blue lines indicate the dsRNA regions. Complete gels are shown in Supplementary Fig. 1C and 1D. (**D**) Toeprinting assay showing the sites of pre-initiation complex formation in the *dnaA* 5’UTR. The reverse transcription step was performed using ^32^P-5’-end-labelled primer A (Supplementary Fig. 2A). Inclusion of 30S ribosomes and tRNA^fMet^ are indicated on the top of the gel. Unfolded 5’UTR lane, assay performed using a heat-unfolded *in vitro* transcribed RNA. U, C, G, A lanes, sequencing reactions using the ^32^P-5’-end-labelled primer A. The large red and orange triangles indicate the reverse transcriptase stops induced by the 30S initiation complexes formed at the AUG and CUG codons, respectively. The AUG and CUG codons are indicated on the sequencing lanes with small red and orange triangles, respectively.

We also assessed the level of sequence conservation by BLAST, using the 5’-most 233 nt of the *dnaA* mRNA of *C. crescentus* NA1000 as a query. Closely related sequences were found in several species of the *Caulobacter* and *Phenylobacter* genera (family *Caulobacteriacea*). Alignment of these 66 sequences revealed three conserved sequence domains. Further analysis, using the co-variance-based software CMfinder, identified three conserved independent structural domains corresponding to helices P1-3, P4, and P6 (Supplementary Fig. 2B), thus additionally supporting the probing results.

### The structure of the *dnaA* 5’UTR inhibits pre-initiation complex formation *in vitro*

One of the most striking features of the proposed *dnaA* mRNA structure is stem-loop P5. Its close proximity to the translation start site might suggest reduced access of initiating ribosomes. Since the *Caulobacter dnaA* 5’UTR had been reported to lack an obvious Shine-Dalgarno (SD) sequence^24^, we first predicted plausible sites of interaction with the 30S ribosomal subunit. The base-pairing energy was calculated for an 8 nt sliding window of the 5’UTR and the anti-SD sequence of *Caulobacter*’s 16S rRNA (Supplementary Fig. 2C), which suggested possible SD-like interactions between A143 and G150.

Experimentally, we assayed translation pre-initiation complex formation *in vitro* by toeprint analysis^25^ on the corresponding *dnaA* mRNA and obtained a weak reverse transcription signal consistent with 30S-tRNA^fMet^ binding at the reported AUG start codon, indicating that the predicted SD (between position 143 and 150) that is buried in stem P5 is functional. Conducting the toeprint after heat-induced denaturation of the RNA gave a significantly enhanced signal. This shows that destabilisation of the proximal elements (i.e., P5, P6) overcomes the partial inhibition seen in Fig. 1D, and suggests that the mRNA adopts a conformation that limits pre-initiation complex formation.

It is noteworthy that our toeprint assay indicated a second band, consistent with complex formation at a non-canonical CUG start codon (nt 114-116), with an upstream SD which was also predicted *in silico* (Fig. 1D and Supplementary Fig. 2C). However, our genetic experiments, as described in detail below, showed that this site is not used for translation initiation *in vivo*.

### A fluorescent reporter system to monitor post-transcriptional regulation of *dnaA* during carbon starvation

To study the *cis*-regulatory elements in *dnaA* mRNA that are involved in the post-transcriptional regulation of DnaA under carbon starvation, we developed an *in vivo* fluorescent reporter system. We translationally fused the *dnaA* promoter and 233 bp of *dnaA* (comprising the 5’UTR and the first 26 codons encoding the DnaA N-terminus, 5’UTR-N_t_ hereafter) to the enhanced green fluorescent protein (eGFP) gene on a low copy plasmid and transformed the construct into wild type *Caulobacter* (Fig. 2A). When shifting the resulting reporter strain from glucose-supplemented to glucose-limiting medium (Fig. 2B, left panel) a transient increase of normalised fluorescence of the culture reached a maximum level, followed by a subtle decline until the end of the experiment. The maximum fluorescence occurred approximately two hours after the culture had entered a starvation-induced growth arrest. Importantly, upon substitution of the *dnaA* 5’UTR-N_t_ with the artificial non-nutritionally controlled 5’UTR_6/13_, eGFP continued to accumulate throughout the starvation phase, while showing similar normalised culture fluorescence during exponential growth as the 5’UTR_6/13_ strain (OD_600_ 0.05 – Fig. 2C, left panel). This result demonstrates that the observed block of eGFP accumulation from two hours after entering growth arrest strongly depends on the presence of the *dnaA* 5’UTR-N_t_.

**Fig. 2.**
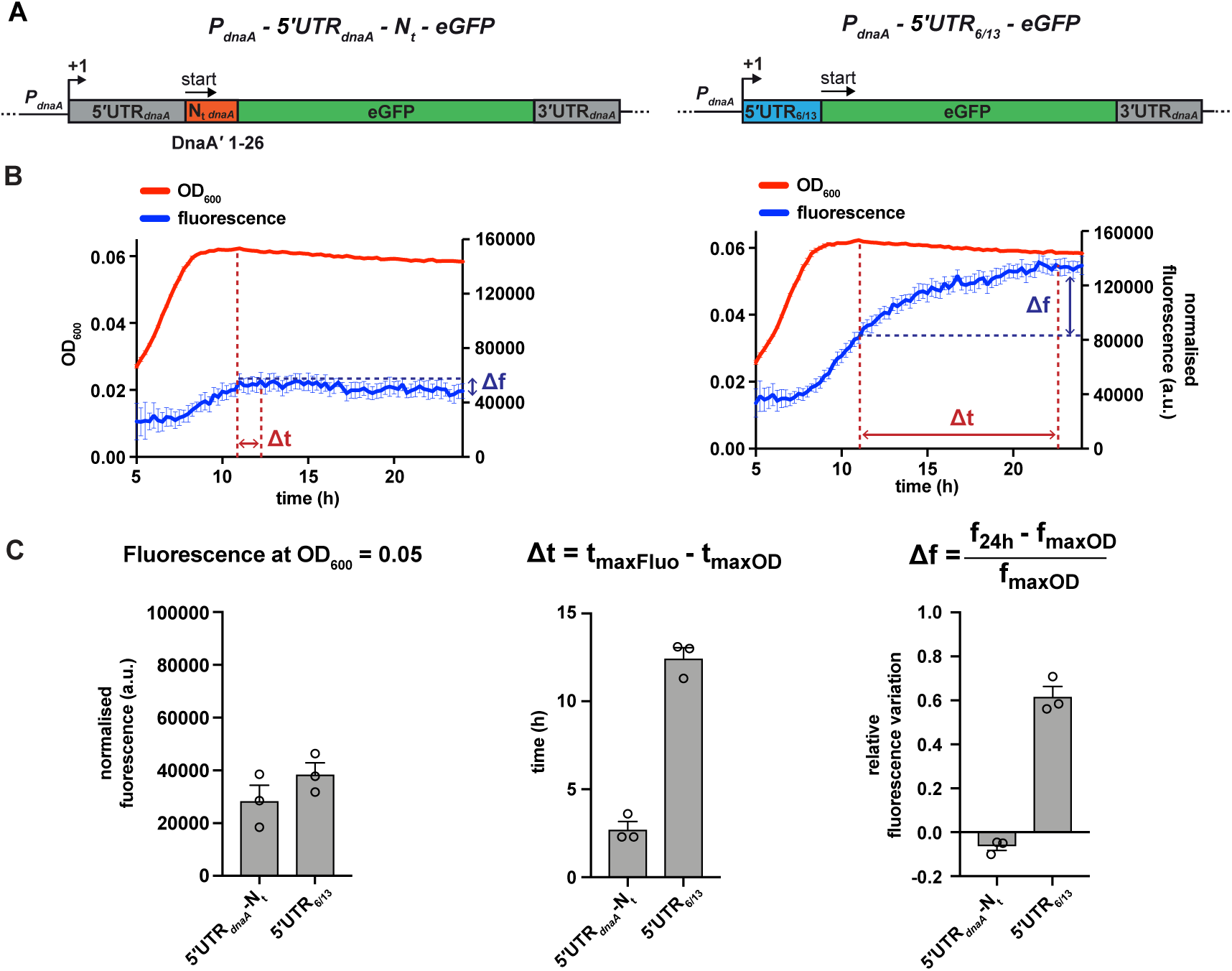
A fluorescent reporter system monitors the post-transcriptional regulation of *dnaA* during carbon starvation. (**A**) Schematic illustration of the plasmid-borne reporter constructs. In the 5’UTR-N_t_ construct (left-hand side), the *dnaA* promoter (*P_dnaA_*), the *dnaA* 5’UTR (5’UTR*_dnaA_* - grey) and the first 26 codons of *dnaA* open reading frame (N_t *dnaA*_ - orange) were translationally fused to the eGFP gene (green), followed by *dnaA* intrinsic terminator (3’UTR*_dnaA_* - grey). In the 5’UTR_6/13_ construct (right-hand side), the 5’UTR*_dnaA_*-N_t *dnaA*_ module was substituted with the non-nutritionally regulated 5’UTR_6/13_ (light blue). (**B**) Culture fluorescence was measured to monitor eGFP synthesis during growth in M2G_1/10_ for 24 hours using a microplate reader. Background correction was performed at each time point by subtracting the fluorescence of a strain carrying a pMR10 empty plasmid. The normalised fluorescence (blue curve) was obtained by dividing with the OD_600_ (red curve) at each time point. (**C**) Values of fluorescence intensity at OD_600_ = 0.05, Δt and Δf calculated using the kinetic profiles in (B). The parameters Δt and Δf are defined by the equations reported on top of the bar graphs and are graphically represented in (B). Averages of three independent replicates are shown with error bars representing standard errors.

To quantitatively describe the different kinetics of eGFP accumulation, we defined two parameters, Δt and Δf. The first expresses the time difference (in hours) between the maxima of the growth and the fluorescence curves (Fig. 2B), providing a measure of the time required to arrest eGFP accumulation in response to carbon exhaustion. The 5’UTR-N_t_ strain gave a much smaller Δt value than the 5’UTR_6/13_ strain (Fig. 2C, middle panel), suggesting efficient downregulation of eGFP expression in response to carbon starvation only when the reporter gene is under the control of the *dnaA* leader. The second parameter, Δf, denotes the relative accumulation of eGFP in the time interval between the culture reaching its maximum OD_600_ and the termination of the measurement (24 h). In this time frame, the 5’UTR_6/13_ strain exhibited a clear increase in fluorescence, resulting in a significantly higher Δf value compared to the 5’UTR-N_t_ strain, which is characterised by a slightly negative Δf value (Fig. 2C, right panel).

### Stem P5 has an inhibitory effect on DnaA translation, but does not mediate DnaA downregulation at the onset of carbon starvation

Having established an *in vivo* reporter system that monitors post-transcriptional regulation of *dnaA* at the onset of carbon starvation, we performed a mutational analysis to identify sequence and structural elements required for the starvation-induced downregulation of DnaA. We first focused on stem P5 and its effect on translation rates in response to nutrient availability. A set of six mutant strains was constructed, with point mutation in the 5’UTR predicted to modify the stability of stem P5 (Fig. 3A). As expected, the stabilising mutation G125C (ΔG_folding_^P^^5^ = ×11.3 kcal/mol) completely abolished eGFP expression, whereas mutations that weaken stem P5 increased eGFP expression during exponential growth (Fig. 3B). In particular, the C127A mutation, predicted to both break one base-pair and entail fraying at the bottom of the stem, caused a strong increase in basal eGFP expression. Most strikingly, none of the five stem P5 mutations affected the pattern of eGFP accumulation during the carbon starvation phase, as reflected by largely unchanged Δt and Δf values compared to the strain with the 5’UTR-N_t_ wild type sequence (Fig. 3C,D and Supplementary Fig. 3A). Therefore, even though stem P5 stability affects *in vivo* translation efficiency *per se*, it is not involved in mediating the nutritional downregulation of DnaA. Additionally, these data suggest that the AGG sequence (nt 143-145), located 10 nt upstream of the AUG start codon, serves as a short and weak SD sequence.

**Fig. 3.**
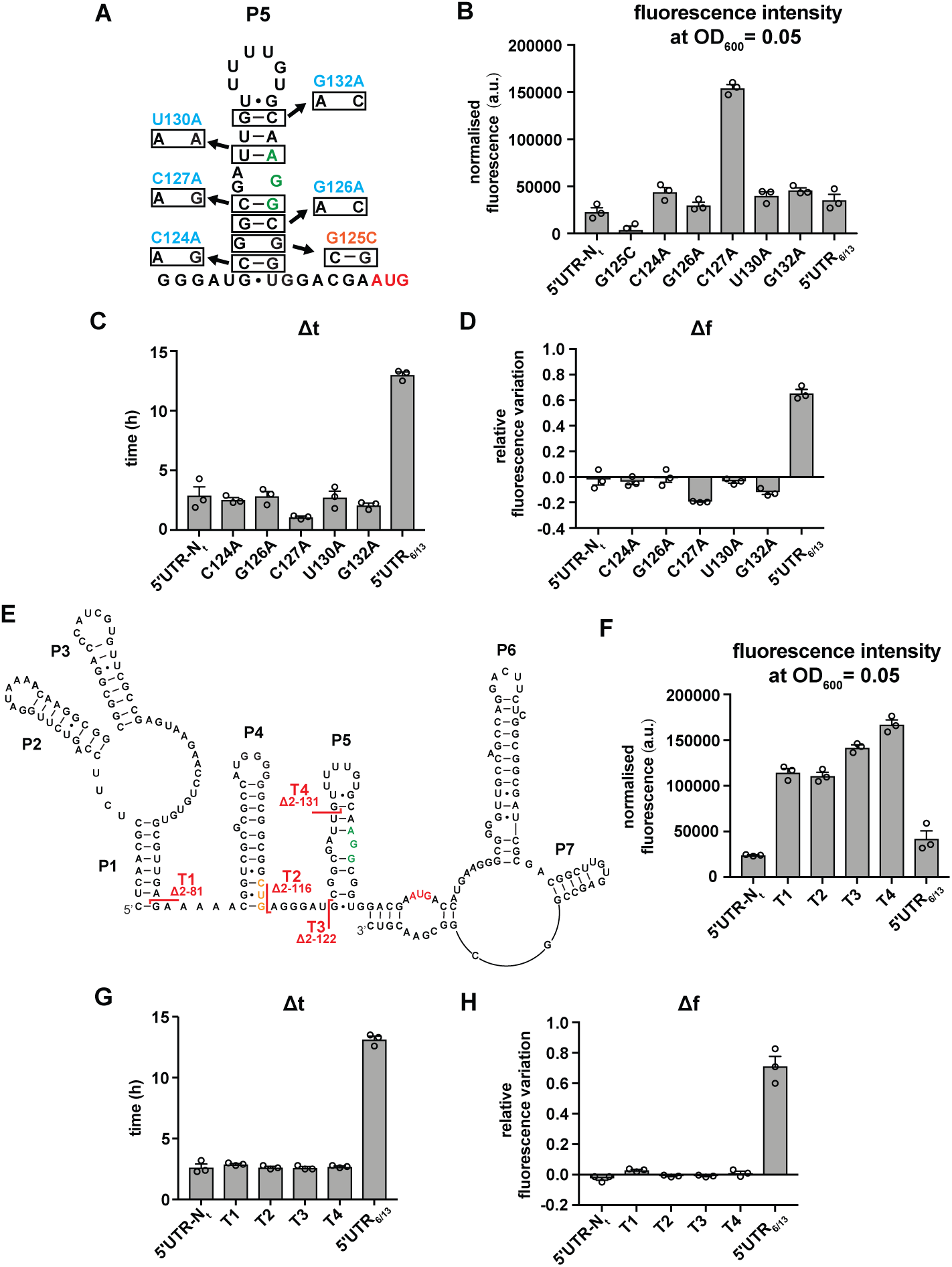
Most of the *dnaA* 5’UTR sequence is not required for the response to carbon starvation. (**A**) Stabilizing (orange) and destabilizing (light blue) mutations introduced in stem P5 by site directed mutagenesis of the 5’UTR-N_t_ reporter construct. The start codon and the putative RBS are indicated in red and green, respectively. (**B-D**) Values of fluorescence intensity at OD_600_ = 0.05, Δt and Δf calculated for the mutants in (A). (**E**) 5’UTR truncation mutants. Sites of truncation (T1-T4) are depicted on the mRNA secondary structure. The start codon and the putative SD are indicated in red and green respectively. The non-canonical CUG codon is coloured in orange. (**F-H**) Values of fluorescence intensity at OD_600_ = 0.05, Δt and Δf calculated for the truncation mutants in (E). See Supplementary Fig. 3A and 3B for growth curves and fluorescence kinetic profiles. All data are shown as averages of three independent replicates with error bars representing standard errors.

To search for other regulatory sequences or structural determinants in the *dnaA* 5’UTR, we engineered a series of truncation mutants in which, starting from the 5’ end of the *dnaA* transcript, increasing portions of the 5’UTR were removed (Fig. 3E). Although truncations T1-T4 increased eGFP expression levels, none of them notably changed Δt or Δf values (Fig. 3F-H and Supplementary Fig. 3B). This result demonstrates that most of the leader is dispensable for the nutritional downregulation of DnaA, and strengthens the conclusion that the putative CUG start codon (see toeprint in Fig. 1D) does not impact DnaA translation *in vivo*. Adding more support, mutation of the canonical AUG start codon or deletion of the entire stem P5 completely abolished DnaA translation, even when the upstream in-frame CUG codon was converted to AUG (Supplementary Fig. 4). Hence, the putative upstream translation start site is not functional *in vivo*.

Two additional deletions (T5, T6) were created to assess whether the loop region of P5 plays a role in regulation of DnaA translation (Supplementary Fig. 5). Truncation T5 showed an eGFP accumulation pattern comparable to truncations T1-T4 and the wild type construct. The T6 mutation removes nearly the entire 5’UTR and corresponds to the ΔUTR*_dnaA_* mutation which, in our earlier study, suggested the *dnaA* 5’UTR to be required for starvation-induced downregulation of DnaA^18^. This mutant gave higher Δt and Δf values compared to the 5’UTR-N_t_ control. However, it also exhibited much lower eGFP levels, indicative of inefficient translation initiation, presumably due to the absence of an extended single-stranded RNA stretch upstream of the AGG SD^27^. Even drastic changes in P5 loop nucleotides (mutants L1-L3) failed to cause substantial changes in Δt and Δf compared to 5’UTR-N_t_ (Supplementary Fig. 5). Thus, the loop region of P5 does not affect regulation of DnaA translation.

In summary, stem P5 limits translation efficiency but does not mediate the starvation-induced downregulation of DnaA. Furthermore, most of the *dnaA* 5’UTR sequence is not required for the response to carbon starvation.

### A sequence element downstream of the *dnaA* start codon mediates nutritional control of translation

Since most putative regulatory elements in the 5’UTR were ruled out as causative for starvation-induced regulation, we considered the mRNA region downstream of the translation start site, which codes for the N-terminal 26 amino acid residues of DnaA (N_t_ hereafter). Strikingly, upon deletion of the N_t_ region (Fig. 4A, 5’UTR-ΔN_t_), eGFP fluorescence continued to accumulate during the carbon starvation phase (Fig. 4B, upper panel). This resulted in significantly increased Δt and Δf values, indicating that this mRNA segment modulates *dnaA* translation at the onset of starvation. The repressing effect of N_t_ was independent of the upstream 5’UTR (Fig. 4B,C); strains with unrelated 5’UTRs (5’UTR_6/13_ or 5’UTR_lac_) showed similar N_t_-dependent changes in the fluorescence kinetic profiles. While deletion of the N_t_ region abolished the nutritional control of DnaA translation, duplication of this region in tandem (2xN_t_) resulted in an earlier response to starvation, as reflected in lower Δt and Δf values compared to the 5’UTR-N_t_ control strain (Fig. 4D,E and Supplementary Fig. 6B). These results demonstrate that the mRNA region downstream of the AUG start codon is required and sufficient for arresting eGFP accumulation during the carbon starvation phase.

**Fig. 4.**
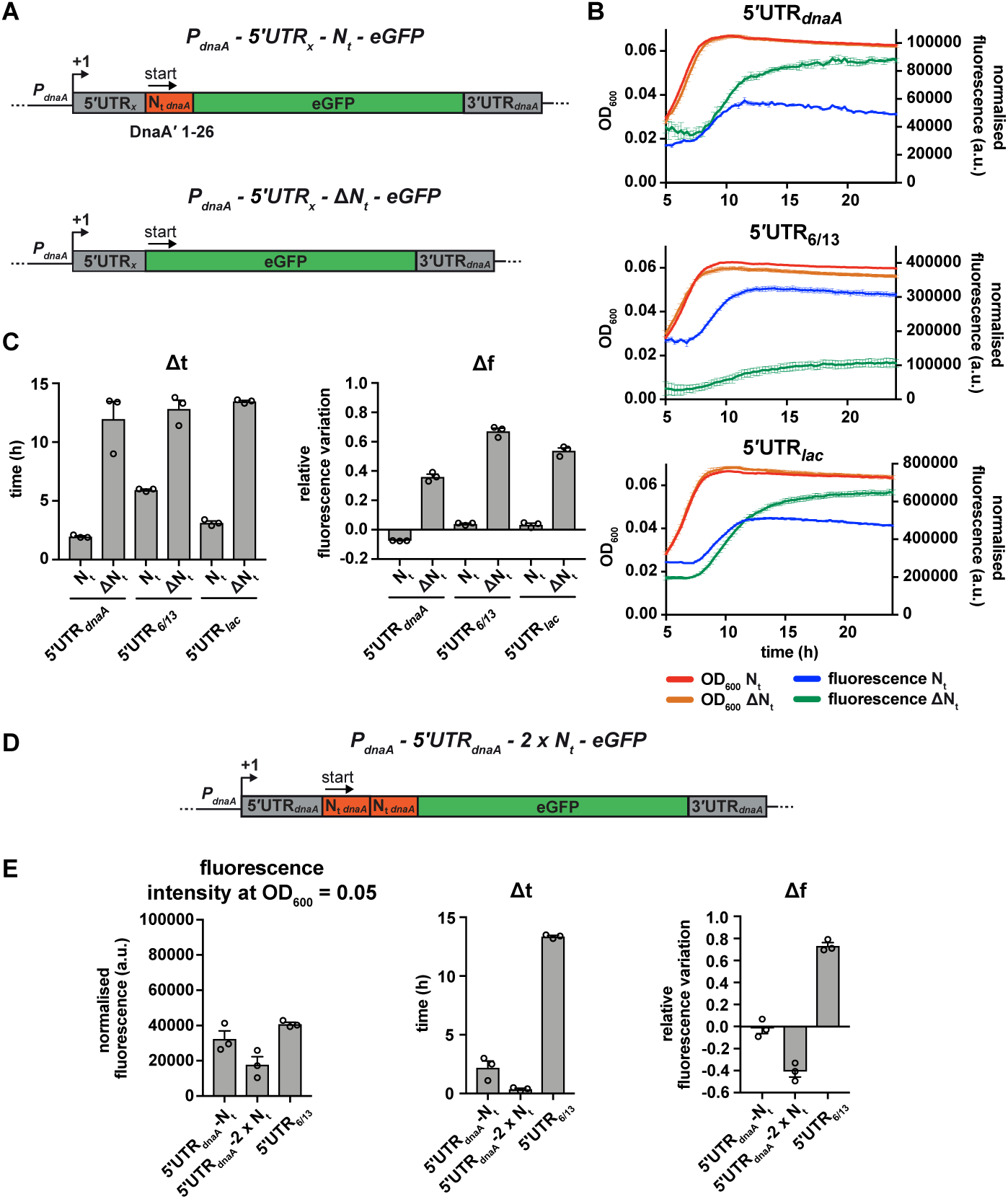
The mRNA region encoding the N-terminus of DnaA mediates the nutritional control of translation independently of the choice of the 5’UTR. (**A**) Schematic illustration of the reporter constructs utilised to study the role of the mRNA region encoding DnaA N_t_ in the translational response to carbon starvation. In the 5’UTR_x_-N_t_ constructs (top) the *dnaA* promoter (*P_dnaA_*) was fused to one of three possible 5’UTRs (5’UTR_x_ = 5’UTR*_dnaA_*, 5’UTR_6/13_, and 5’UTR*_lac_* – grey) followed by the first 26 codons of *dnaA* open reading frame (N_t *dnaA*_ - orange) and the eGFP gene (green). In the 5’UTR_x_-ΔN_t_ constructs (bottom) the N_t_ module is absent. (**B**) Growth curves and fluorescence kinetic profiles of the 5’UTR_x_-N_t_ (red and blue) and 5’UTR_x_-ΔN_t_ (orange and green) constructs. (**C**) Δt and Δf values of the 5’UTR_x_-N_t_ and 5’UTR_x_-ΔN_t_ reporter constructs. (**D**) The 5’UTR*_dnaA_*-2xN_t_ reporter construct was obtained by duplicating the N_t_ module (orange). (**E**) Values of fluorescence intensity at OD_600_ = 0.05, Δt and Δf calculated for the 2xN_t_ strain. The relative growth curves and fluorescence kinetic profiles are shown in Supplementary Fig. 6B. All data are shown as averages of three independent replicates with error bars representing standard errors.

Because the N_t_ region is translationally fused to eGFP, the stability of the reporter protein might be affected. Therefore, we compared eGFP protein stability between the 5’UTR-N_t_ and 5’UTR-ΔN_t_ strains at the onset of starvation, after chloramphenicol-induced translation shut-down followed by Western blotting and plate reader-based measurements of fluorescence. Both assays showed that the eGFP reporter was stable in both strains (Supplementary Fig. 7), excluding that the presence of *dnaA* N_t_ at the N-terminus of eGFP significantly affects protein stability. This result is also congruent with a previous study which showed that the N-terminus of DnaA is required but not sufficient for DnaA proteolysis^28^. Additionally, the plate reader experiment showed that the increase in fluorescence in the 5’UTR-ΔN_t_ strain during the carbon starvation phase was completely abolished in the presence of chloramphenicol (Supplementary Fig. 7B,C), confirming that the fluorescence increase in this strain depends on ongoing translation during starvation. Consistent with our previous results (Fig. 4B,C), this effect was independent of the choice of 5’UTR.

### Downregulation of DnaA translation occurs during the early elongation phase and requires N_t_’s native reading frame

We considered two alternatives by which N_t_ might influence DnaA translation. Either the secondary structure of the mRNA segment encoding N_t_ (i.e., P6 and P7) inhibits translation^29, 30^, or the nature of the codons or amino acids encoded by this sequence affects translation elongation. To discriminate between these possibilities, we constructed an N_t_ double frameshift mutant (dfsN_t_) that entirely changes the codon and peptide sequence between residues 3 and 26 (Fig. 5A and Supplementary Table 1) while preserving the mRNA secondary structure, as determined by mFOLD. Strikingly, similar to the 5’UTR-ΔN_t_ strain, this mutant continued to synthesize eGFP also during the carbon starvation phase, reflected in high Δt and Δf values compared to the 5’UTR-N_t_ strain (Fig. 5B,5C). This result argues against a role of secondary structure elements present in N_t_ mRNA and instead suggests that the maintenance of the native reading frame is essential for nutritional control of translation.

**Fig. 5.**
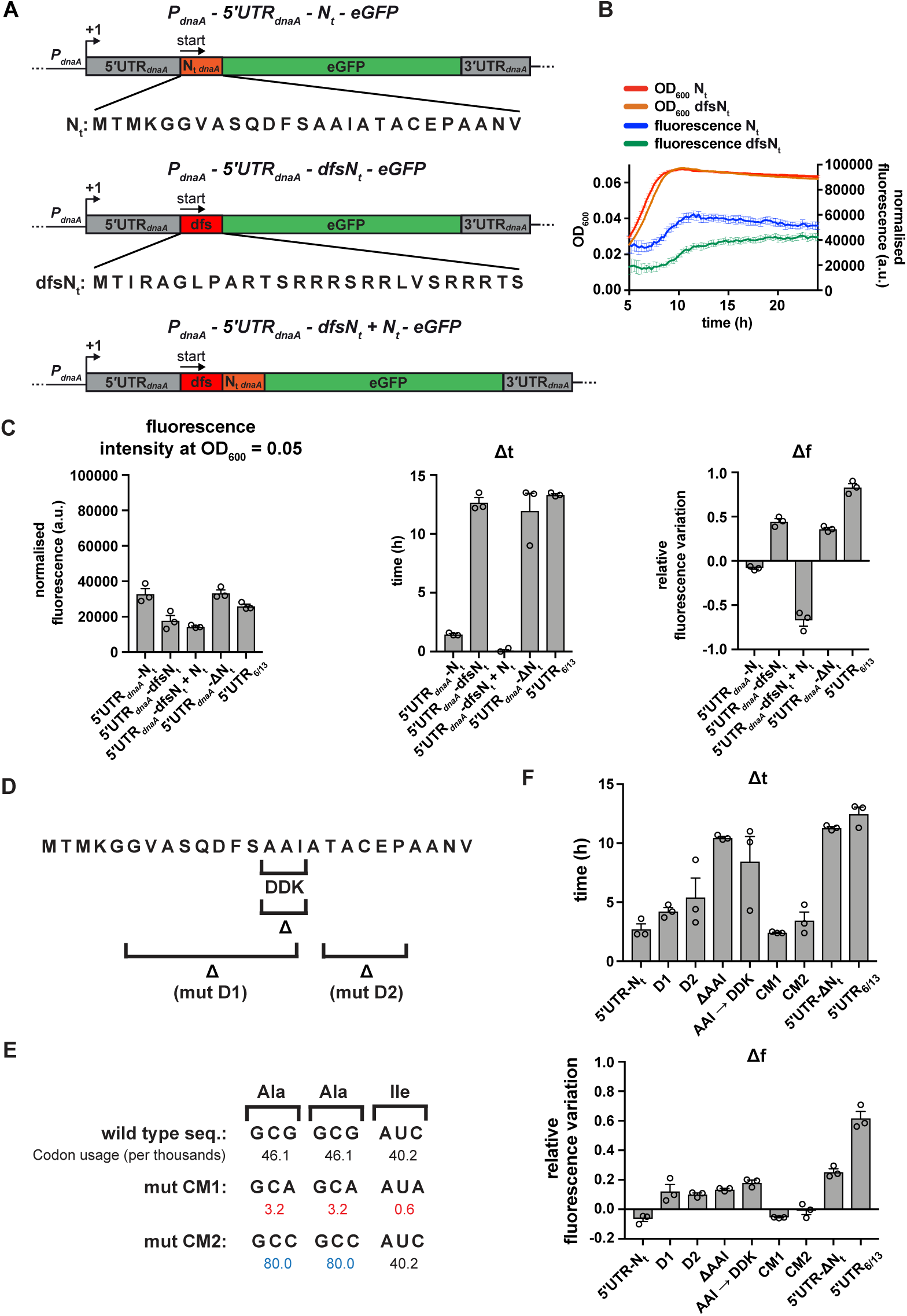
A specific amino acid sequence motif in the N-terminus of DnaA inhibits translation elongation under carbon starvation. (**A**) Schematic illustration of the double frame-shift mutant reporter constructs. Top, wild type amino acid sequence of DnaA N_t_ in the context of the 5’UTR*_dnaA_*-N_t_ reporter construct. The first double frame-shift mutant (5’UTR_dnaA_-dfsN_t_ – in the middle) was generated upon deletion of G164 and insertion of a G after residue C233 by site-directed mutagenesis (Supplementary Table 1). The double frame-shift mutation alters the reading frame of N_t_ (dfsN_t_ module, in red), without affecting the eGFP frame. The 5’UTR*_dnaA_*-dfsN_t_ + N_t_ construct (bottom) was generated by introducing the aforementioned double frame-shift mutation in the first N_t_ module of the 2xN_t_ construct (Fig. 4D). (**B**) Comparison of fluorescence kinetic profiles between the 5’UTR-N_t_ (blue) and the dfsN_t_ (green) reporter strains. (**C**) Values of fluorescence intensity at OD_600_ = 0.05, Δt and Δf calculated for the reporter strains in (A). (**D**) Illustration of deletions D1, D2 and ΔAAI, and the amino acid substitution AAI–DDK. (**E**) Illustration of the AAI motif codon mutations CM1 and CM2. Top, the wild type codon sequence encoding the AAI motif, along with the values of codon usage. In mutant CM1 and CM2, the AAI motif is encoded by codons that present either lower (values in red) or higher (values in blue) usage. (**F**) Values of Δt and Δf calculated for the mutant reporter strains illustrated in (D) and (E). Data are reported as averages of three independent replicates with error bars representing standard errors.

We next tested whether the inhibitory effect of N_t_ was maintained when this region was moved further downstream within the *dnaA* coding region, or depended on its close proximity to the translation start site. For this, we made use of the tandem 2xN_t_ strain and disrupted the first N_t_ region by introducing the two frameshift mutations to generate a dfsN_t_-N_t_ sequence (Fig. 5A). Despite a reduction in basal eGFP expression, this strain showed clearly reduced Δt and Δf values (Fig. 5C and Supplementary Fig. 6C). This indicates starvation-dependent downregulation of *dnaA* and suggests that the N_t_ sequence inhibits translation elongation even when it is located further downstream within the open reading frame.

In conclusion, our data are compatible with a model in which DnaA translation is downregulated under carbon starvation during the early stages of polypeptide chain elongation due to the specific amino acid or codon composition of N_t_.

### A specific nascent chain amino acid sequence inhibits translation elongation under starvation

Having established that the N_t_-mediated downregulation of DnaA translation depends on the native reading frame, we next wanted to identify sequence features in N_t_ that are critical for regulation. A BLAST revealed high conservation of the DnaA N_t_ amino acid sequence in several members of the *Caulobacteriacea* family (Supplementary Fig. 6A). A general feature of the identified sequences is a high degree of hydrophobicity, with alanine being the most abundant amino acid. Using our fluorescent reporter, we designed two mutants in which regions of DnaA N_t_ were deleted; mutant D1 lacks a hydrophobic region spanning residues 6 to 15, and mutant D2 lacks residues 18 to 22 (Fig. 5D). Both D1 and D2 resulted in moderately higher Δt and Δf values (Fig. 5F and Supplementary Fig. 6D) compared to the 5’UTR-N_t_ strain, indicating a delay in starvation-dependent inhibition of DnaA translation. However, none of these mutations recapitulated the phenotype observed in the ΔN_t_ or dfsN_t_ strains. In order to identify shorter motifs responsible for the phenotype observed in the D1 mutant, we generated additional amino acid deletions and substitutions in this segment of DnaA N_t_. Most of these did not affect eGFP accumulation during carbon starvation (Supplementary Fig. 8). However, the deletion or substitution of one group of three residues (A14-A15-I16; AAI motif hereafter) in the conserved hydrophobic region significantly increased both Δt and Δf values compared to the 5’UTR-N_t_ control, similar to the 5’UTR-ΔN_t_ and 5’UTR-dfsN_t_ strains (Fig. 5F and Supplementary Fig. 6D). These results indicate that the amino acids of the AAI motif contribute to the inhibition of DnaA translation.

Previous work established that synonymous codons, decoded by distinct isoacceptor aminoacyl (aa)-tRNAs (i.e., charged with the same amino acid), can display differences in their sensitivity to starvation^9, 31^. For instance, the charged levels of major aa-tRNAs, which read the most abundant codons, rapidly decrease upon amino acid starvation^31^. However, when the AAI motif codons were mutated to either less or more abundant synonymous codons (Fig. 5E), no significant effects on the starvation-dependent inhibition of DnaA translation were observed (Fig. 5F and Supplementary Fig. 6D). This suggests that the amino acid sequence of the N-terminus of DnaA mediates the starvation-induced translation arrest irrespective of codon choice. It is of note that the introduction of less abundant codons, containing A or T in the third position, resulted in higher eGFP expression levels (Supplementary Fig. 6D), probably due to a decreased propensity for a stable mRNA structure near the translation start site^29, 30, 32^.

Thus, our data suggest that, under carbon-limiting conditions, translation elongation efficiency decreases in a sequence-dependent manner during the synthesis of the DnaA N-terminus. Importantly, this effect depends on the amino acid composition of the DnaA N_t_, but not the codons encoding it.

### The sequence-specific translation downregulation mechanism operates in a heterologous host

We wondered whether the identified starvation-responsive sequence element also can function in a different cellular context, and chose to analyse the effect of the N_t_ region of the *Caulobacter dnaA* mRNA in *Escherichia coli*. For this, we transcriptionally fused the 5’UTR-N_t_-eGFP or 5’UTR-ΔN_t_-eGFP modules, respectively, from the *Caulobacter* reporter to an IPTG-inducible Lambda O1 promoter on a low copy plasmid, and transformed the resulting construct into *E. coli* (Fig. 6A). Indeed, shifting to carbon-limiting conditions, eGFP accumulation ceased completely in the strain with the *Caulobacter* N_t_ sequence, while it continuously increased fluorescence in the strain lacking this region (Fig. 6B). A similar repressing effect was also observed when the 5’UTR of *Caulobacter dnaA* was replaced by the artificial 5’UTR_6/13_, demonstrating that N_t_ is sufficient for regulating translation in response to carbon starvation also in *E. coli*. These data show that the identified starvation-responsive sequence encoded within the N-terminus of DnaA constitutes an autonomous regulatory element that can operate in a heterologous host.

**Fig. 6.**
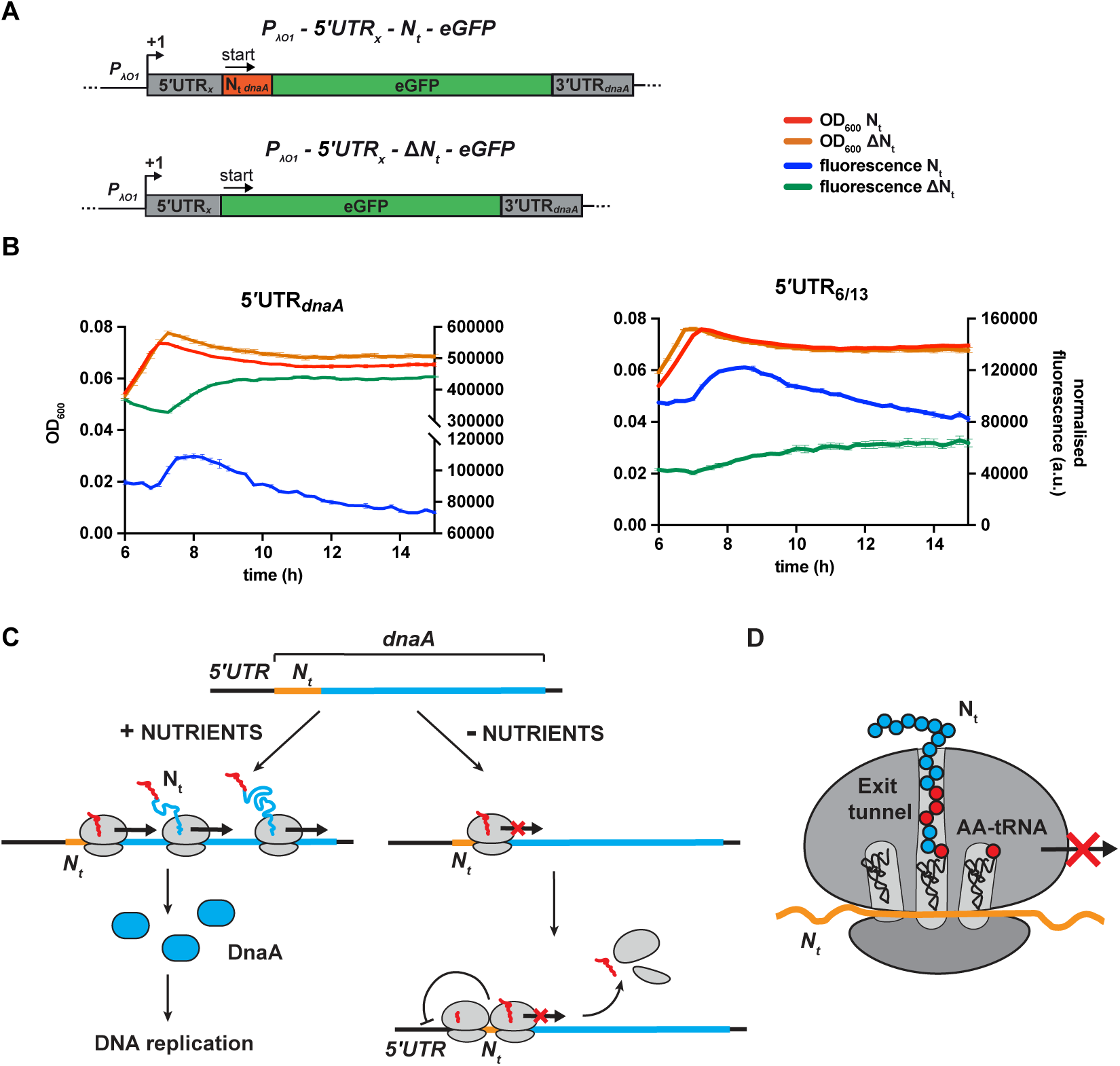
Starvation-induced inhibition of DnaA translation elongation by an autonomously operating nascent chain sequence element. (**A**) Schematic illustration of the reporter constructs utilised to study DnaA N_t_ functionality in the heterologous host *E. coli*. In the 5’UTR_x_-N_t_ constructs (top) the IPTG-inducible Lambda promoter O1 (*P_λO1_*) was fused either to the 5’UTR*_dnaA_* or the 5’UTR_6/13_ (5’UTR_x_ - grey) followed by *Caulobacter*’s N_t_ (N_t *dnaA*_ - orange) and *eGFP* (green). Similar to Fig. 4A, the 5’UTR_x_-ΔN_t_ constructs (below) lack the N_t_ module. The constructs were cloned into the low-copy plasmid pZE12 and transformed into *E. coli* MG1655. (**B**) The *E. coli* reporter strains were grown for 15 hours in M9 medium supplemented with 0.02% glucose in a microplate reader. Normalised culture fluorescence was measured to monitor eGFP synthesis in the reporter strains (5’UTR*_dnaA_*-N_t_ versus 5’UTR*_dnaA_*-ΔN_t_ on the left-hand side and 5’UTR_6/13_-N_t_ versus 5’UTR_6/13_-ΔN_t_ on the right-hand side). Background subtraction and fluorescence normalisation were performed as described for the *Caulobacter* reporter strains. Data are reported as averages of three independent replicates with error bars representing standard errors. (**C**) The proposed model for the regulation *dnaA* translation under carbon starvation. In the presence of nutrients (left-hand side), *dnaA* mRNA is efficiently translated leading to accumulation of active DnaA (light blue) that, in turn, triggers DNA replication initiation. Under carbon starvation (right-hand side), translation rate decreases during the first stages of the elongation phase at a specific sequence located downstream of the translation start site (*N_t_* –orange). The resulting ribosome traffic jams possibly stimulate disassembly of the leading ribosome from the mRNA and/or inhibit new cycles of translation initiation. The regulatory nascent polypeptide (N_t_) is shown in red. (**D**) The downregulation of *dnaA* translation requires specific amino acid residues (in red) encoded in the region downstream of the start codon (*N_t_* – in orange). These specific amino acids arrest translation after or while being added to the growing DnaA nascent polypeptide (N_t_).

## DISCUSSION

The integration of nutritional information with essential cellular processes such as protein synthesis and DNA replication is critical for cellular life, but poorly understood on a molecular level. This study reveals a new mechanism, by which some bacteria can regulate the synthesis of the replication initiator DnaA in response to nutrient availability by modulating the rate of translation. In the model organism *C. crescentus*, we identified a specific N-terminal sequence in the nascent DnaA polypeptide chain that acts as a novel regulatory element which tunes DnaA translation elongation in response to the cellular nutrient and growth status. With its crucial function in bacterial replication initiation, the abundance and activity of DnaA must be precisely regulated in response to environmental and cellular changes to ensure cellular survival.

### A model for dnaA regulation under carbon starvation

Based on our results, we propose a model of nutritionally controlled DnaA translation. Under optimal conditions, efficient transcription of *dnaA* and a high rate of DnaA translation ensure the accumulation of active DnaA that triggers DNA replication initiation (Fig. 6C). At the onset of carbon starvation, translation ceases during the early stage of the elongation phase at a specific sequence located downstream of the *dnaA* start codon (as discussed below). Ribosome pausing at this sequence is expected to result in ribosome traffic jams that, as previously proposed, stimulate premature translation termination either directly, by inducing disassembly of the leading ribosome from the mRNA, or indirectly, by recruiting ribosome rescue systems^9, 33^. Due to the proximity of the stalling site and the start codon, persistent queues of 2-3 ribosomes might also decrease translation rates by physically preventing access of initiating ribosomes to the translation start site^34–36^. Additionally, ribosome pausing may result in increased mRNA decay or Rho-dependent termination of transcription^37^. The postulated starvation-dependent elongation pause significantly reduces the rate of DnaA synthesis. The concomitant degradation of full-length DnaA by Lon then aids the rapid clearance of DnaA from the cell^18^, preventing DNA replication initiation.

### The molecular mechanism of starvation-induced translation inhibition

The nutritional downregulation of DnaA translation depends on a specific sequence downstream of the *dnaA* start codon. Since neither the RNA secondary structure of this sequence nor the codon choice impacted the nutritional regulation of DnaA (Fig. 5), it seems unlikely that the RNA structure spatially hinders ribosome movement, or that the starvation-dependent reduction of a specific set of isoacceptor aa-tRNAs causes ribosome stalling. Instead, we suggest that the amino acids encoded by this sequence arrest translation after, or while, being added to the growing DnaA polypeptide (Fig. 6D). This finding is in line with the recent observation that the identity of the amino acids encoded by early codons of a gene can impact protein synthesis yield^38^. Previous work has also demonstrated that certain nascent peptide sequences specifically arrest translation through interactions with the ribosome exit tunnel surface, thereby regulating the expression of downstream genes through translational coupling, or affecting transcription termination and mRNA decay^39, 40^. A well characterised example is the force-sensing arrest peptide SecM, which contributes to the translational control of the downstream gene *secA* that encodes a translocase subunit^41, 42^. Another class of arrest peptides modulates gene expression by interacting with small effector molecules in the ribosome exit tunnel^43, 44^. The AAI motif identified in this study, possibly along with neighbouring amino acids in the N-terminus of DnaA, might interact with the ribosome in a similar way as arrest peptides. However, while arrest peptides commonly serve as *cis*-acting elements that regulate the expression of downstream genes, the sequence element that we describe represents, to our knowledge, the first example of a regulatory nascent polypeptide sequence in the N-terminus of a protein that directly modulates translation efficiency of the same protein in response to nutrient availability.

### Possible ways of sensing nutrients

The identified nascent sequence element specifically decreases overall DnaA translation rates at the onset of carbon starvation, demonstrating that it is responsive to the nutrient status of the cell. Since this is recapitulated in a heterologous host (Fig. 6B), it argues for a sensing mechanism that is triggered by a global starvation-induced change in translation strategy^8^, rather than by *Caulobacter-*specific accessory factors. For example, reduction in the supply of one or more components of the translation machinery (e.g., ribosomes, aa-tRNAs, elongation factors, GTP) could in principle slow down overall translation rates^8, 37, 45^. Consequently, translating ribosomes could become more susceptible to starvation-specific inhibitory sequences akin to the sequence identified here. Alternatively, a highly conserved *trans*-acting factor, a metabolite or a signalling molecule could be involved in the nutrient sensing mechanism. Such a putative factor or small molecule might either release the translational arrest during nutrient-rich conditions or enhance it during starvation.

### The dnaA 5’UTR as a regulatory element

This work presents the first detailed structural and functional characterisation of the *dnaA* 5’UTR. Although the 5’UTR is not required for modulating translation rates under starvation, it might play other regulatory roles. Consistent with a previous study^46^, our results show the inhibitory effect of the *dnaA* leader on translation *per se*, and demonstrate that the efficiency of translation initiation can be significantly increased by reducing the stability of the secondary structure around the translation start site. Interestingly, although the *dnaA* 5’UTR, like the majority of mRNAs in *Caulobacter*, lacks an obvious SD sequence^24^, our results suggest that a short AGG motif buried in stem P5 could function as an SD. The modulation of SD accessibility by a *trans*-acting factor (e.g., a regulatory RNA or protein) interfering with stem P5 structure could conceivably be a strategy to directly regulate the initiation of *dnaA* translation in response to specific external or internal cues. Consistent with this idea, a previous computational study proposed the *dnaA* 5’UTR of *Caulobacter* to be a possible regulatory target of multiple small non-coding RNAs^47^.

In conclusion, we have identified a new sequence-dependent mechanism that modulates translation elongation of a specific protein in response to nutrient availability, highlighting the importance of the first stages of translation elongation as a regulatory target. It is particularly striking that the mechanism described functions to control the concentration of the highly conserved replication initiator DnaA. This protein is critical for DNA replication in nearly all bacteria, and nutritional control mechanisms that modulate DnaA synthesis contribute to the correct timing of DNA replication initiation in response to changes in nutrient availability and growth rate. The finding that DnaA in *Caulobacter* and related bacteria is regulated at the level of translation elongation reveals important new layers of control acting on this important protein.

The conclusions reported here may suggest a more general pattern. We consider it possible that the proposed mechanism of translational downregulation via nascent peptide sequence-dependent effects potentially tunes the expression of other genes during starvation. Principally, this mechanism could help to globally remodel the proteome under starvation conditions when cells need to reallocate their resources from growth-promoting to maintenance functions.

## AUTHOR CONTRIBUTIONS

M.F. performed the experiments and carried out the computational and data analysis. K.J. and M.F. conceived the study and wrote the manuscript with input from E.G.H.W and C.R.. All authors contributed to the design of the experiments and the interpretation of the results.

## DECLARATION OF INTERESTS

The authors declare no competing interests.

## ACKNOWLEDGMENTS

We thank members of the Jonas lab for helpful discussions and comments on the manuscript and Dr. Iker Irisarri for bioinformatic advice. The study was financially supported by the European Union’s Horizon 2020 research and innovation program under the Marie Skłodowska-Curie grant agreement No 797801, the Swedish Foundation for Strategic Research (FFL15-0005), the Swedish Research Council (K.J.: 2016-03300, E.G.H.W.: 2017-03765), and funding from the Strategic Research Area (SFO) program distributed through Stockholm University.

## MATERIALS AND METHODS

### Growth conditions

All *Caulobacter* reporter strains were derived from the wild-type strain NA1000 (Supplementary Table 2), and were grown routinely in PYE or M2G (minimal medium with 0.2% glucose) at 30°C while shaking at 200 rpm. For the carbon exhaustion experiments, exponentially growing cells were washed and inoculated in M2G_1/10_ (0.02% glucose). *E. coli* DH5*α* was used as a host for plasmid construction and was routinely grown in LB medium at 37°C while shaking at 200 rpm. The *E. coli* reporter strains were derived from wild-type strain MG1655 (Supplementary Table 2). Pre-cultures were grown at 30°C while shaking at 200 rpm in LB or M9 minimal medium supplemented with 0.2% glucose. Carbon exhaustion experiments in the plate reader were performed in M9 medium supplemented with 0.02% glucose and 1 mM IPTG. When appropriate, the following antibiotic concentrations were used: 1 μg/mL kanamycin (PYE and M2G liquid media), 25 μg/mL kanamycin (PYE plates), 30μg/mL kanamycin (LB liquid medium), 50 μg/mL kanamycin (LB plates), 50 μg/mL ampicillin (LB liquid medium) and 100 μg/mL ampicillin (LB plates).

### Bacterial strains and plasmid construction

All the plasmids in this study were constructed by Gibson assembly^48^. pMR10-BG (Supplementary Fig. 9A) was derived from pMR10^49^ by exchanging *lacZα* and *oriT* with a multicloning site. The vector backbone for the pMR10-5’UTR-N_t_-GFP (Supplementary Fig. 9B) construct was obtained upon digestion of pMR10-BG with HindIII and BamHI, while the insert was first assembled in a smaller vector, pMCS^50^, and then amplified by PCR with appropriate primers. The insert comprised the following sequences, in order, (i) the *rrnB1*-T1T2 terminator, (ii) 245 bp upstream of the *dnaA* transcription start site (*P_dnaA_*), (iii) the 5’UTR*_dnaA_*, (iv) the first 78 bp of the *dnaA* open reading frame, (v) the eGFP gene (fused in-frame) and (vi) 63 bp downstream of the *dnaA* stop codon (3’UTR*_dnaA_*). The other pMR10 reporter constructs (Supplementary Table 1) were generated by site-directed mutagenesis using pMR10-5’UTR-Nt-GFP as template. All constructs were finally transformed into *C. crescentus* NA1000 by electroporation.

Plasmid pZE12-BG (Supplementary Fig. 9C) was generated from pZE12-luc plasmid^51^ by deleting the luciferase gene and part of the *P_λO1_* promoter. The pZE12 reporter plasmids were obtained by replacing the luciferase gene in the pZE12-luc plasmid, with the 5’UTR_x_-N_t_-eGFP or 5’UTR_x_-ΔN_t_-eGFP modules amplified from the related pMR10 constructs (Supplementary Fig. 9D). All pZE12 constructs were transformed into *E. coli* MG1655 as previously described^52^.

### Immunoblotting

Pelleted cells were resuspended in 1x SDS sample buffer (1/10 volume of *β*-mercaptoethanol added before use), normalised to the optical density of the culture (40 μL per units of 0.1 OD_600_), and heated to 95°C for 10 min. Protein extracts were subjected to SDS-PAGE for 90 min at 130 V at room temperature on 12% Mini-PROTEAN TGX Stain-Free gels (Bio-Rad #4568046). Samples were transferred to nitrocellulose membranes using a BioRad Trans-Blot Turbo system (“standard sd” protocol). To verify equal loading, total protein was visualised using the 2,2,2-trichloroethanol in-gel method^53^ after PAGE separation and prior to blotting. DnaA and eGFP were detected with anti-DnaA^54^ (1:5,000 dilution) and anti-GFP (Thermo Fisher Scientific #A11122, 1:20,000 dilution) primary antibodies, respectively, followed by a 1:5,000 dilution of goat anti-rabbit secondary horseradish peroxidase-conjugated antibody (Thermo Fisher Scientific #31460). After treating the membranes with the SuperSignal Femto West reagent (Thermo Fisher Scientific #34094), blots were scanned with a LI-COR Odyssey Fc imaging system. Fiji (ImageJ) was used for image processing and band intensity quantification.

### *In vitro* transcription of RNA

A DNA template containing a T7 promoter was generated by PCR using *Caulobacter crescentus* NA1000 genomic DNA as template and the following primers: 5’-GAAAT<u>TAATACGACTCACTATAG</u>CTCAACGCTCTTCCAGTCTTGG-3’ (Forward, T7 promoter underlined) and 5’-CACCCAGCTCACGCTTCAAAG-3’ (Reverse). The Megascript Kit (Life Technologies, #AM1330) was used for *in vitro* transcription. The RNA was purified on a 6% denaturing polyacrylamide gel. The band corresponding to the transcribed RNA was detected by UV-shadowing and cut from the gel. The RNA was then eluted into 300 mM Na-acetate, 0.1% SDS, and 1 mM EDTA. After phenol–chloroform–isoamyl alcohol (25:24:1) extraction and ethanol precipitation, the RNA pellet was dissolved in water. RNA concentration was measured by NanoDrop and the quality was assessed by denaturing PAGE. 5’-end-labeling of CIAP-treated RNA (Invitrogen, #18009-019) with T4 PNK (Thermo Fisher Scientific, #EK0031) in buffer A and [*γ*-P^32^]-ATP (10 mCi/mL, 3,000 Ci/mmol-Perkin Elmer # BLU002A). The labelled RNA was gel-purified as described above.

### Native gel electrophoresis

The *in vitro* transcribed RNA was refolded for 1 min at 95°C, followed by 1 min on ice and 10 min at 30°C in 1x TMK buffer (100 mM K-acetate, 5 mM Mg-acetate, 50 mM Tris-HCl pH 7.5). The sample (about 10,000 cpm) was directly loaded on a 6% non-denaturing polyacrylamide gel. The gel was run at 4°C for 2 h (200 V) in 0.5x TEB, then covered with plastic film, and exposed to a storage phosphor screen for 1 h. Signals were detected using a PMI system (Bio-Rad) and analysed by Image Lab 6.1 (Bio-Rad).

### Structural Probing

Two probing approaches were used to determine the secondary structure of the *in vitro* transcribed RNA. In the first, 0.1 pmol of ^32^P-5’-end-labelled RNA were subjected to RNase T1 or Pb^2+^ probing at 30°C in the presence of 2 μg carrier yeast tRNA (Invitrogen #AM7119) (Supplementary Fig. 1B). The labelled RNA was first denatured for 1 min at 95°C, followed by 1 min on ice. Before adding the carrier tRNA and the probes, the labelled RNA was allowed to re-fold for 10 min in a buffer containing 10 mM Tris-acetate pH 7.5, 100mM K-acetate, 5 mM, DTT and 10 mM Mg-acetate (10 μL total reaction volume). RNase T1 probing was done using 1 unit of enzyme (Invitrogen, #AM2283) for 5 min, and stopped by adding 40 μL cold 0.3 M Na-acetate. Chemical probing was done with Pb^2+^-acetate (Sigma-Aldrich-Merk) at a final concentration of 10 mM and 25 mM for 5 min. Reactions were stopped with addition of cold EDTA (50 mM final concentration) followed by 35 μL of cold 0.3 M Na-acetate. After phenol–chloroform–isoamyl alcohol extraction, RNA was ethanol-precipitated. RNA pellets were dried and dissolved in loading dye. About 10,000 cpm of each sample were loaded on a 12% denaturing polyacrylamide gel, fixed for 5 min (10% ethanol, 6% acetic acid), and transferred to 3-mm Whatman paper. Signals were detected using a storage phosphor screen and a PMI system (Bio-Rad), and analysed using Image Lab 6.1 (Bio-Rad). Cleavage pattern induced by the probes were assigned using RNase T1 hydrolysis under denaturing conditions (i.e., T1 ladder) and an alkaline ladder. The ladders were prepared following the manufacturer’s instructions (RNase T1 manual, Invitrogen). We used 0.2 pmol of 5’-end-labelled RNA/ 2 μg of carrier yeast tRNA for the T1 ladder, and 0.1 pmol of 5’-end-labelled RNA/ 1 μg of carrier yeast tRNA were used for the alkaline ladder.

The second probing approach used primer extension to detect cleavage positions after treatment with RNases T1 and V1 (Supplementary Fig. 1C,D). Here, reactions were performed as described above, but using 1 pmol of unlabelled RNA and 1 μg carrier yeast tRNA. RNase V1 probing was done using 0.1, 0.02, or 0.01 enzyme units (Invitrogen, #AM2275) for 5 min, stopped by adding 40 μL of cold 0.3 M Na-acetate. T1 ladders were done in presence of 1 pmol of unlabelled, denatured RNA and 2 μg of carrier yeast tRNA. The probed RNA and the ladders were phenol–chloroform–isoamyl alcohol extracted and precipitated. Dried pellets were then resuspended in water, and primer extension was performed at 37°C for 20 min using 100 U of SuperScript IV reverse transcriptase (Invitrogen, #18090010) and two different ^32^P-5’-end-labelled primers: 5’-CACCCAGCTCACGCTTCAAAG-3’ (primer A) or 5’-GAGAAGTCCTGGCTGGCAAC-3’ (primer B). Primers were labelled using T4 PNK in buffer A and [*γ*-P^32^]-ATP (10 mCi/mL, 3,000 Ci/mmol) and purified using G-50 Microspin columns (GE Healthcare, #27-5330-01). Reverse transcription reactions were stopped by adding 40 μL of cold 0.3 M Na-acetate. After phenol/chloroform extraction, the RNA template was removed by addition of 3 M KOH for 3 min at 90°C, followed by 1 hour incubation at 37°C. After base neutralisation with HCl, the cDNAs were ethanol-precipitated, centrifuged, dissolved in loading buffer, and resolved on 12% polyacrylamide gels. Gels were fixed and treated as above. The sequencing reactions were done using the USB Thermo Sequenase Cycle Sequencing Kit (Affymetrix #78500) and ^32^P-5’-end-labelled primers A and B. The template for the sequencing reaction was obtained by PCR using *Caulobacter crescentus* NA1000 genomic DNA and the following primers: 5’-CAAATATTTACAAAGGCCGATCAGGG-3’ (Forward) and primer A (Reverse).

### Toeprinting assay

The toeprinting assay was performed in a 10 μL reaction volume using 0.2 μM of unlabelled RNA and, when present, 100 nM of *E. coli* 30S subunits and 300 nM of *E. coli* initiator tRNA^fMet^ ^55^. The reaction buffer contained 10 mM Tris-acetate pH 7.6, 100 mM K-acetate, 1 mM DTT, and 10 mM Mg-acetate. The RNA was denatured for 1 min at 90°C in the presence of 5’-end-labelled primer A (3 μL, about 50,000 cpm/μL) and 0.5 mM dNTPs. After 1 min on ice, Mg-acetate was added. RNAs were refolded for 5 min, followed by addition of activated 30S (10 min at 30°C). Next, initiator tRNA^fMet^ was added, and 30S-initiation complexes were allowed to form for 20 min. In the unfolded RNA sample, the 30S subunits and the initiator tRNA^fMet^ were added immediately after the 1-min incubation at 90°C. Reverse transcription was started by addition of 50 U of SuperScript IV for 20 min, and stopped by adding 40 μL of cold 0.3 M Na-acetate. The reverse transcription reactions were treated as above and resolved on a 12% denaturing polyacrylamide gel.

### Homologous sequence search and alignments

Three complementary approaches were used to identify homologous sequences of *Caulobacter*’s UTR*_dnaA_* and N_t *dnaA*_. In the first one, an input nucleotide sequence including both the 5’UTR and the first 25 codons of DnaA was used in a BLASTn search against the NCBI nucleotide collection^56^ and the genome database (March 2020), using an E-value threshold of 10^-6^. In the second approach, aiming to identify more distant homologs, the *Caulobacter* DnaA protein sequence (CCNA_00008) was used in a BLASTp search against the annotated proteins in NCBI’s representative bacterial genome database (E-value cut-off 10^-6^). The resulting hits were aligned with mafft v7.427^57^, and a maximum likelihood phylogenetic tree built with IQTREE v. 1.6.10^58^ in order to select the cluster of the most closely related sequences for further analysis. Finally, a tBLASTn search was performed against the NCBI database of representative bacterial genomes, to identify additional unannotated homologs. As previously, a tree was inferred on the hits and only sequences forming a cluster of highly related sequences were analysed further. For each hit resulting from the BLASTp and the tBLASTn searches, the upstream region containing the 5’UTR was extracted from the corresponding genome using the coordinates of the BLAST results. The results of the three homology searches were manually screened, pooled, and aligned using the MUSCLE alignment tool^59^ as implemented in SeaView 5.0.4^60^.

### Computational analysis

mFold^61^ and the RNAfold server of Vienna RNA^62^ were used for RNA secondary structure predictions at 30°C. For a search of putative SD sequences in the 5’UTR of *dnaA* mRNA, we used the RNAcofold tool of the Vienna RNA package to calculate the ΔG of interaction at 30°C between an 8 nt sliding window of the 5’UTR and the anti-SD sequence of *Caulobacter*’s 16S rRNA. Three different anti-SD sequences were considered: 5’-CCUCC-3’, 5’-CCUCCU-3’ and 5’-CACCUCCU-3’. The structural analysis based on co-variance was performed using CMfinder 0.4.1.4^63^ according to the developer’s indications.

### Reporter assays

A Tecan Spark multimode microplate reader was used for culture incubations as well as OD_600_ and eGFP fluorescence measurements. Reporter strains were grown in 150 μL of medium at 30°C in sterile 96-well transparent plates with flat-bottom and lid (Greiner Bio-One # 655182). Initial culture OD_600_ was 0.02 (1 cm optical path length). The plates were orbitally shaken at 180 rpm with a 3 mm amplitude using a humidity cassette to protect from evaporation. OD at 600 nm and fluorescence were measured every 15 min. The following instrument settings were used to measure eGFP fluorescence: excitation wavelength = 485 nm (20 nm bandwidth), emission wavelength = 535 nm (25 nm bandwidth), detector gain = 90 (manual), and automatic z-position optimisation.

Raw data were processed similarly to Zaslaver et al.^64^ using R and the splines package. The OD_600_ measurements of the cultures were background-subtracted using the average value of the blank wells at each time point. To estimate the value of the background fluorescence, we grew a strain carrying the empty pMR10-BG plasmid in each plate. By plotting fluorescence against OD_600_, we noticed that background changed non-linearly over the growth curve. To account for this variation, we fitted the pMR10-BG fluorescence and OD_600_ data with a smoothing spline (“smooth.spline”), generating a mathematical function that allows calculation of the expected background fluorescence at any given OD_600_. Background correction was performed by subtracting the expected background fluorescence to the reporter strain’s fluorescence at each time point. Finally, the resulting value was normalised by dividing it with the OD_600_.

Δt and Δf were calculated as indicated in Fig. 2. The times of maximum fluorescence (t_maxFluo_) and maximum OD_600_ (t_maxOD_), as well as the fluorescence values at 24 hours (f_24h_) and at maximum OD_600_ (f_maxOD_) were calculated after fitting growth and fluorescence curves with smoothing splines (“smooth.spline”). To determine the fluorescence intensity at OD_600_ = 0.05, we first determined the time at which OD_600_ equals 0.05 using “interSpline” and “backSpline” functions. Then, the obtained value was used to calculate the corresponding fluorescence with the R function “predict”. All statistical analyses and graph preparations were performed in GraphPad Prism (version 7).

### Translation shut-off assays

For the Western blot-based translation shut-off assay, cells were cultured in 25 mL of M2G_1/10_ medium for 7 h until growth arrested. Translation was shut off by addition of chloramphenicol (100 µg/mL final concentration). One mL aliquots were withdrawn at 15 min intervals and snap-frozen in liquid nitrogen before being analysed by Western blotting.

For the eGFP translation shut-off assay in the plate reader, reporter strains were grown on M2G_1/10_ for 9 h as above. When appropriate, translation was shut off by adding 100 µg/mL of chloramphenicol to the culture. Fluorescence and OD_600_ were measured at 10 min intervals for 8 h. Plate reader data processing was performed as described above.

---

**Supplementary Fig. 1.**
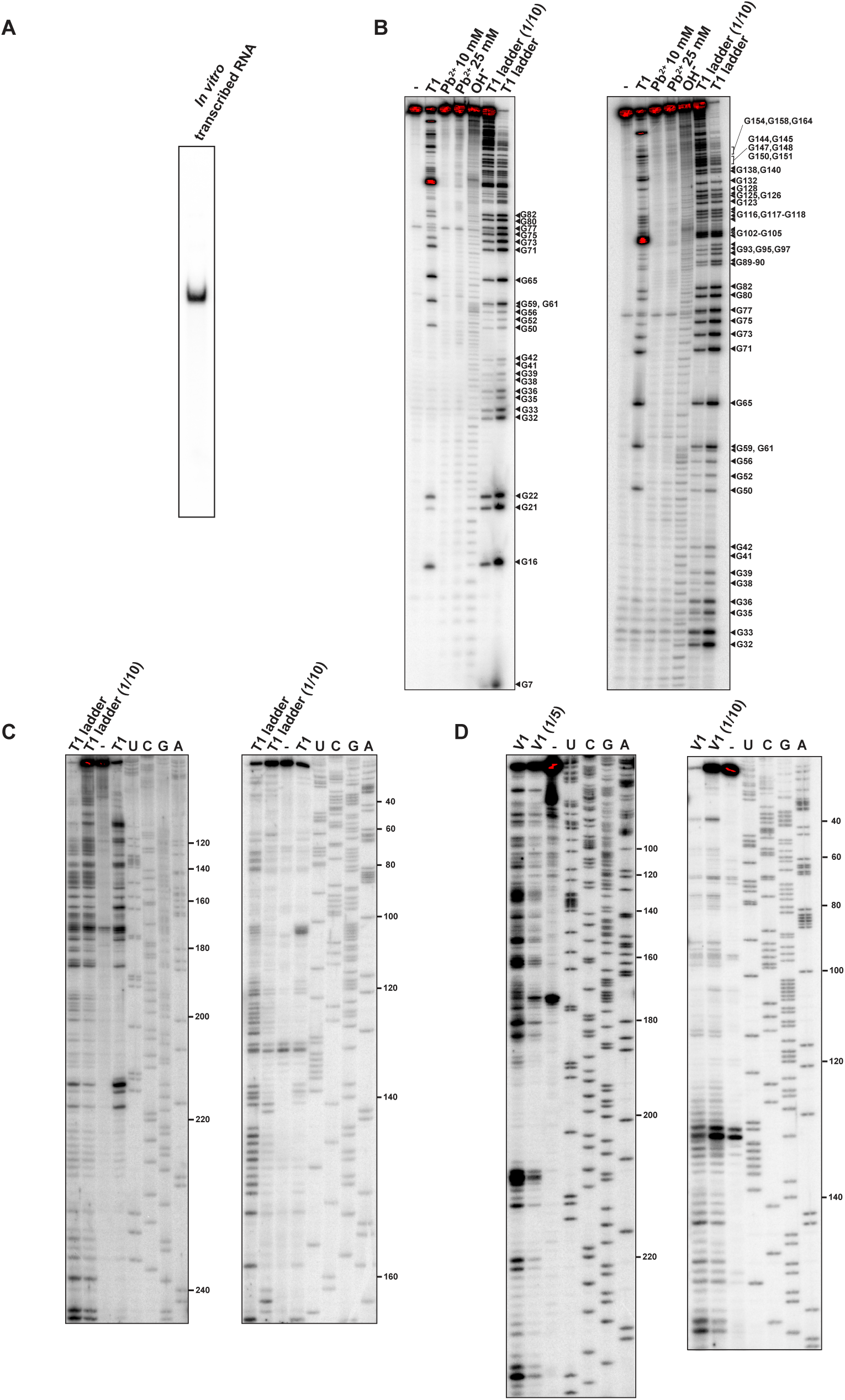
Structural characterisation of the *dnaA* mRNA leader. (**A**) Native 6% PAGE of the *in vitro* transcribed ^32^P-5’-end-labelled RNA comprising the 5’UTR and the first 118 nt of the *dnaA* coding region (total RNA length 273 nt). (**B**) The ^32^P-5’-end-labelled RNA was subjected to RNase T1 and Pb^2+^ structure probing, as described in STAR Methods. Cleavage products were separated on a 12% denaturing polyacrylamide gel and detected by phosphorimager. -, mock-treated RNA. OH^-^, alkaline ladder. T1 ladder and T1 ladder (1/10) lanes, RNase T1 treatment under denaturing conditions using different concentrations of the probe. The gel on the left was run for 2 h, and the one on the right for 4 h. (**C-D**) RNase T1 and V1 structure probing gels obtained using primer extension. The reverse transcription step was performed with two different ^32^P 5’-end-labelled primers (Supplementary Fig. 2A): primer A (left gel) or primer B (right gel). Reverse transcription products were separated on a 12% sequencing gels and detected by phosphorimager. Lanes representing T1 or V1 treatment are indicated, with diluted enzyme concentrations shown (e.g. 1/5 etc.), and U, C, G, A are sequencing reactions using the same primers. -, mock-treated RNA. The probing data (B-D) gave structural information from positions 7-234.

**Supplementary Fig. 2.**
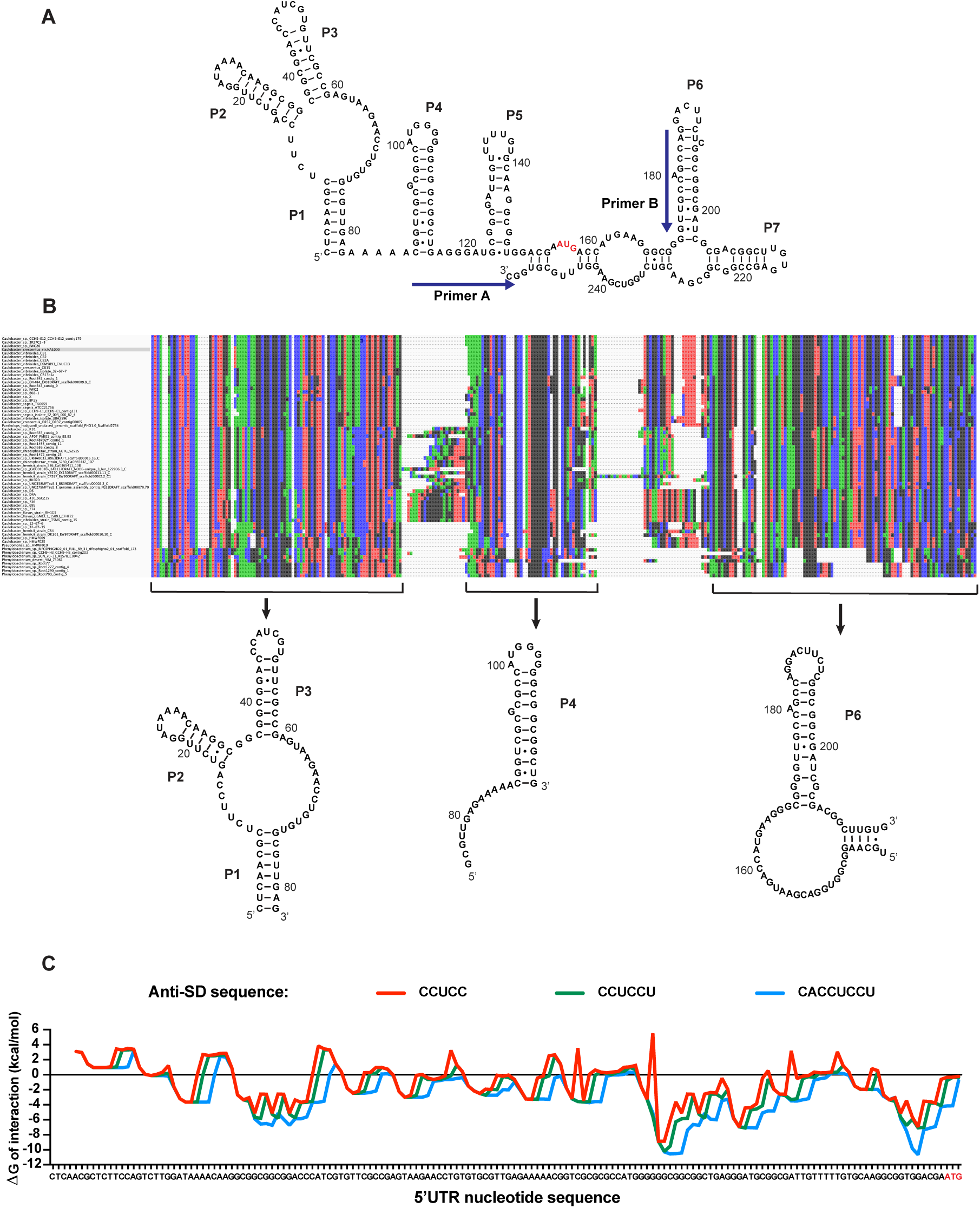
Structure conservation of the *dnaA* mRNA leader and predicted interaction with Caulobacter’s 16S rRNA. (**A**) Alternative model for the secondary structure of the mRNA region encompassing the 5’UTR and the first 98 nucleotides of the *dnaA* open reading frame. The AUG start codon is highlighted in red. Primers A and B, used for probing and toeprinting assays, are represented on the structure. (**B**) Alignment of the 66 sequences identified in the *Caulobacteriacea* family (MUSCLE). The BLAST hits were used to search for conserved structural elements using CMfinder. The co-variance-based software predicted the existence of 3 conserved structural domains corresponding to the helixes P1-P3, P4 and P6. (**C**) Energy of interaction between an 8 nt sliding window of the 5’UTR and the anti-SD sequence of *Caulobacter*’s 16S rRNA. The ΔG of interaction was calculated using the RNAcofold tool of the ViennaRNA package. Three different anti-SD sequences were considered: CCUCC (red), CCUCCU (green) and CACCUCCU (blue).

**Supplementary Fig. 3.**
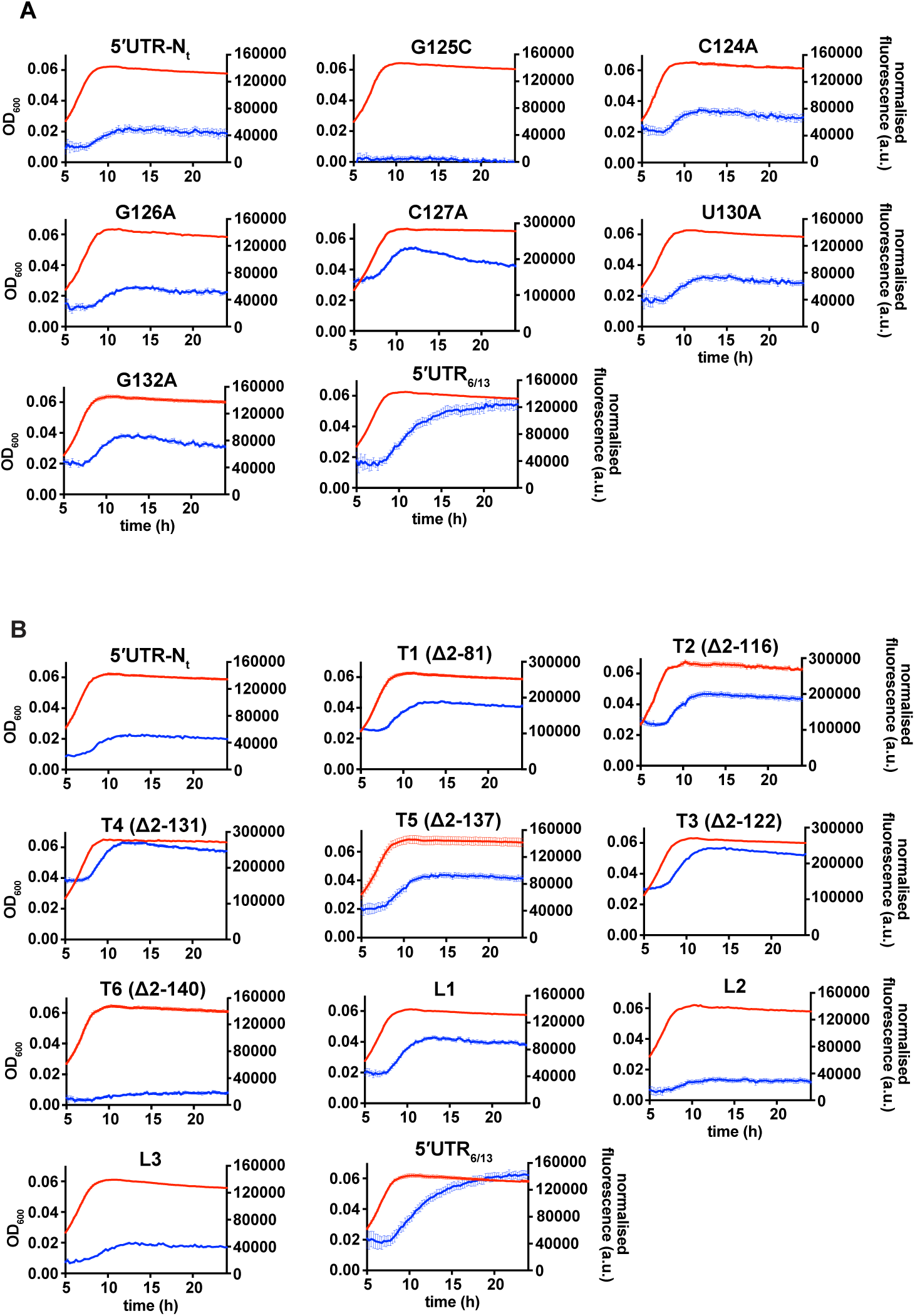
Fluorescence kinetic profiles and growth curves of the reporter strains carrying mutations in the 5’UTR of the dnaA mRNA. (**A**) Reporter strains carrying mutations that stabilize (G125C) or destabilize (C124A, G126A, C127A, U130A, G132A) stem P5. (**B**) Reporter strains with 5’UTR truncations (T1-T6, Fig. 3E and Supplementary Fig. 5A) or loop P5 mutations (L1-L3, Supplementary Fig. 5A). The profiles of the 5’UTR-N_t_ and 5’UTR_6/13_ strains are shown for comparison. Averages of three independent replicates are shown with error bars representing standard errors.

**Supplementary Fig. 4.**
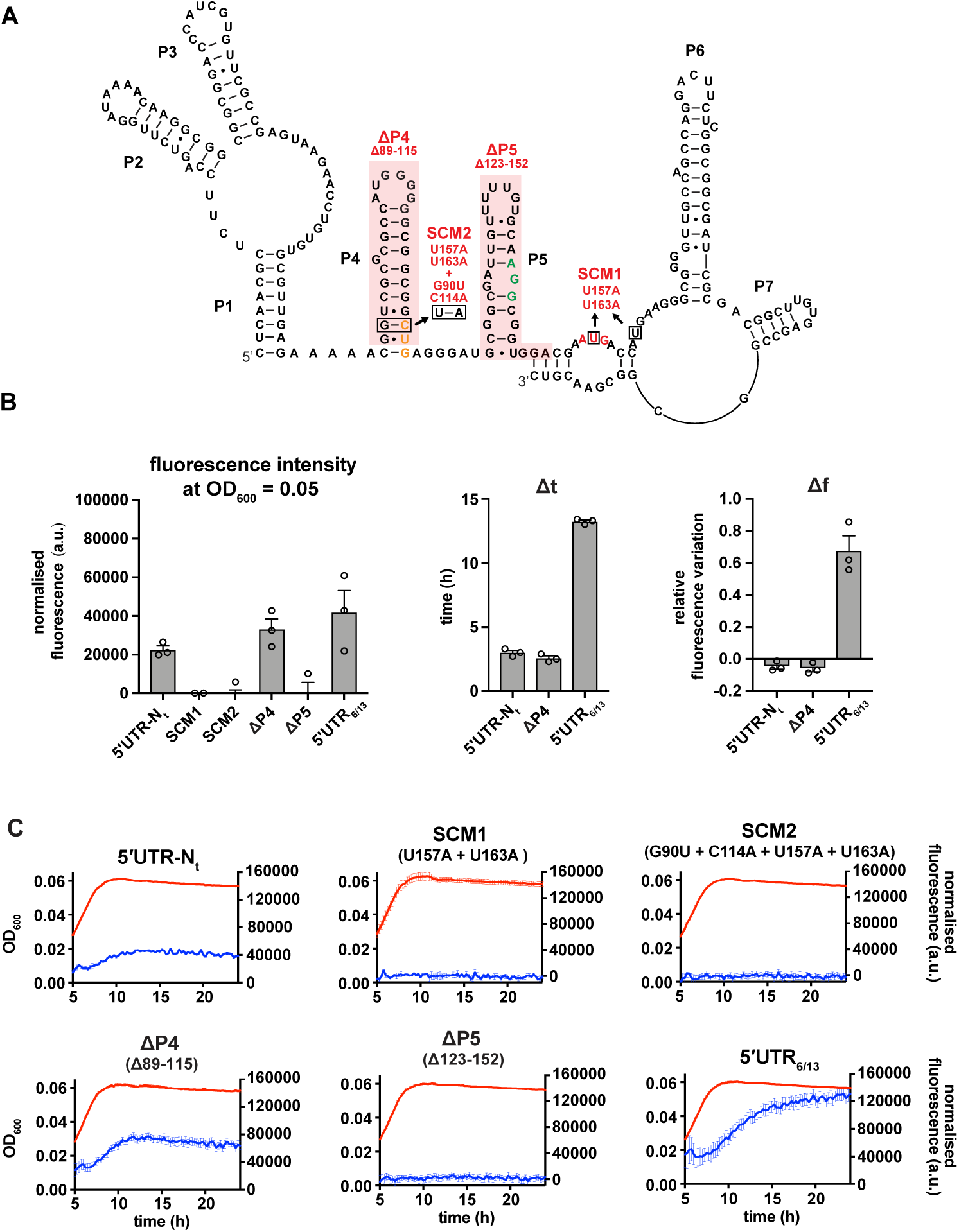
The reported AUG start codon is the only functional translation start site *in vivo*. (**A**) Start codon mutant 1 (SCM1) disrupts the canonical AUG start site (red). In the double mutant SCM2, the upstream CUG codon (orange) is additionally mutated to AUG. ΔP4 and ΔP5: deletions of secondary elements P4 and P5 as indicated (shaded in red). The ΔP5 mutation preserves the frame between the CUG and AUG codons. The putative SD is shown in green. (**B**) Values of fluorescence intensity at OD_600_ = 0.05, Δt and Δf calculated for the reporter mutant strains in (A). (**C**) Growth curves and fluorescence kinetic profiles of the reporter mutant strains in (A). Averages of three independent replicates are shown with error bars representing standard errors.

**Supplementary Fig. 5.**
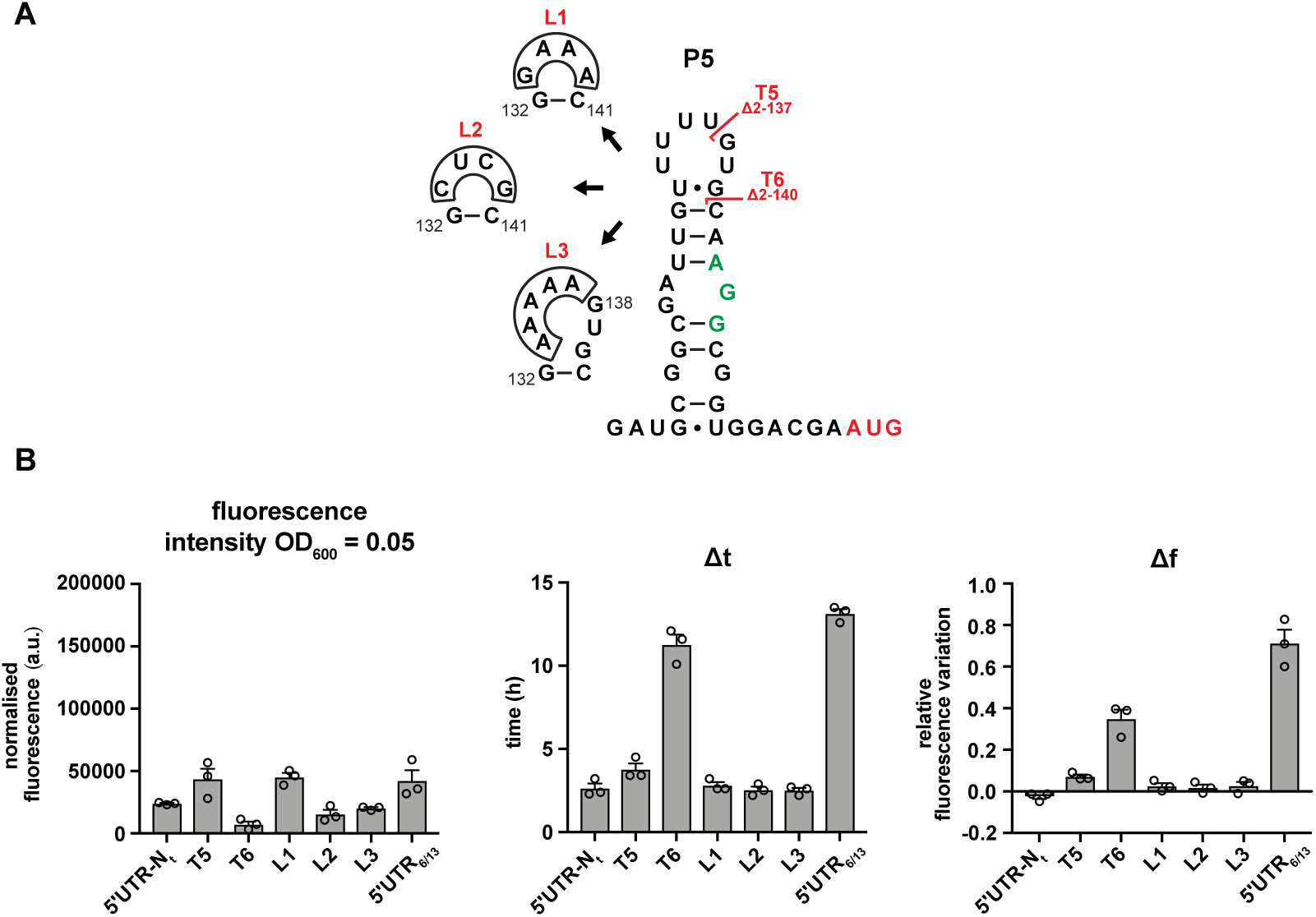
The loop region of P5 does not have a role in DnaA translation regulation. (**A**) Schematics of T5 and T6 5’UTR truncation mutants, and the loop P5 mutations (L1-L3). The start codon and the putative SD are coloured in red and green respectively. (**D**) Values of fluorescence intensity at OD_600_ = 0.05, Δt and Δf calculated for the mutant strains in (A). For the fluorescence kinetic profiles, see Supplementary Fig. 3B. Averages of three independent replicates are shown with error bars representing standard errors.

**Supplementary Fig. 6.**
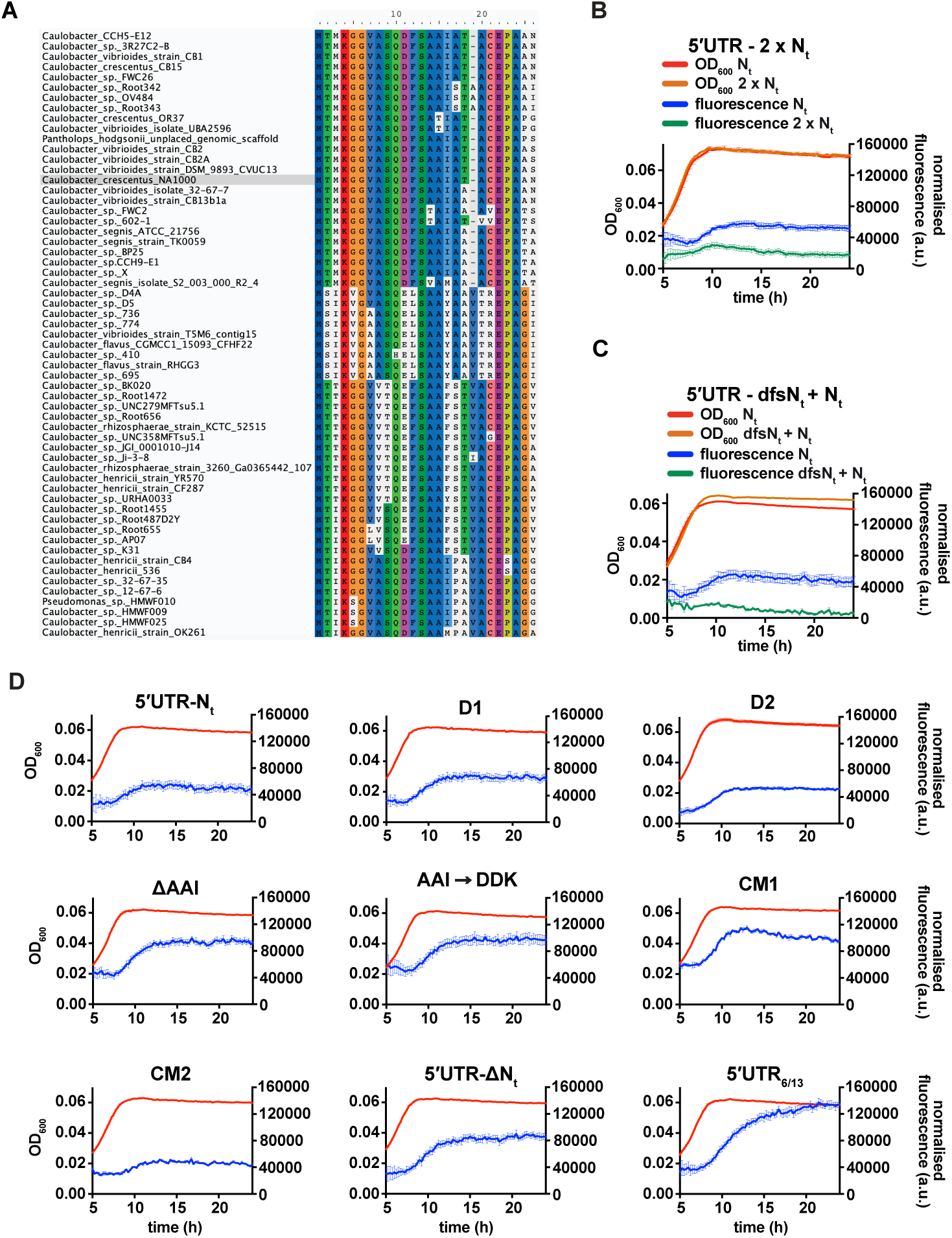
Fluorescence kinetic profiles and growth curves of the reporter strains carrying mutations in DnaA N_t_. (**A**) Amino acid sequence conservation of DnaA N_t_. 59 homologous sequences from the *Caulobacteriacea* family were aligned using MUSCLE. *Caulobacter crescentus* NA1000 sequence is highlighted in grey. (**B**) Comparison of fluorescence kinetic profiles between the 5’UTR-N_t_ (blue) and the 2xN_t_ (green) reporter strains (See Fig. 4D, E). (**C**) Comparison of fluorescence kinetic profiles between the 5’UTR-N_t_ and the dfsN_t_ + N_t_ reporter strains (See Fig. 5A, C) (**D**) Fluorescence kinetic profiles of the reporter strains carrying the deletions D1, D2, ΔAAI, the amino acid substitution AAI-DDK and the codon mutations CM1 and CM2 (See Fig. 5D-F). The profiles of the 5’UTR-N_t_, 5’UTR-ΔN_t_ and 5’UTR_6/13_ strains are shown for comparison. Averages of three independent replicates are shown with error bars representing standard errors.

**Supplementary Fig. 7.**
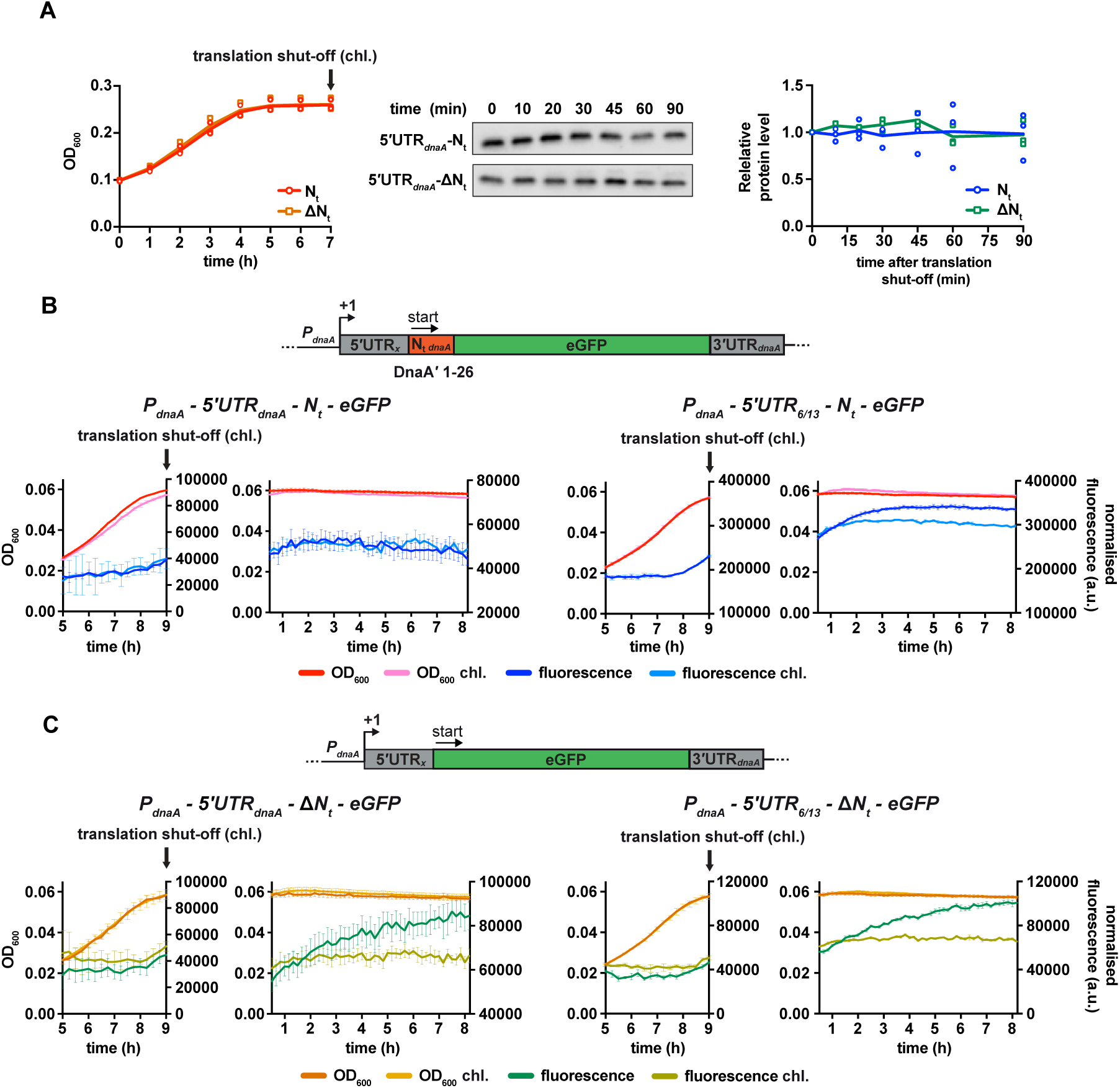
The presence of DnaA N_t_ at the N-terminus of eGFP does not change protein stability. (**A**) Western blot-based translation shut-off assay. The 5’UTR*_dnaA_*-N_t_ and 5’UTR*_dnaA_*-ΔN_t_ reporter strains were cultured in M2G_1/10_ medium for 7 h until growth arrested (left panel). Translation was shut off by adding chloramphenicol (100 µg/mL), and the decrease in eGFP abundance was monitored for 90 minutes by Western blot (middle panel). Right panel: quantification of band intensities. Averages of three independent replicates are shown with error bars representing standard errors. (**B-C**) eGFP translation shut-off assay using fluorescence as a read-out of protein abundance. Reporter strains were grown in M2G_1/10_ in a plate reader for 9 h (i.e., onset of carbon starvation). Translation was stopped by adding chloramphenicol (100 µg/mL) to the culture. Bulk culture fluorescence was measured at regular intervals over 8 h. (B) Fluorescence kinetic profiles of the 5’UTR_x_-N_t_ strains (5’UTR_x_ = 5’UTR*_dnaA_* or 5’UTR_6/13_) in the presence (light blue) or in the absence (dark blue) of chloramphenicol treatment. (C) Fluorescence kinetic profiles of the 5’UTR_x_-ΔN_t_ strains (5’UTR_x_ = 5’UTR*_dnaA_* or 5’UTR_6/13_) in the presence (light green) or absence (dark green) of chloramphenicol treatment. Averages of three independent replicates are shown with error bars representing standard errors.

**Supplementary Fig. 8.**
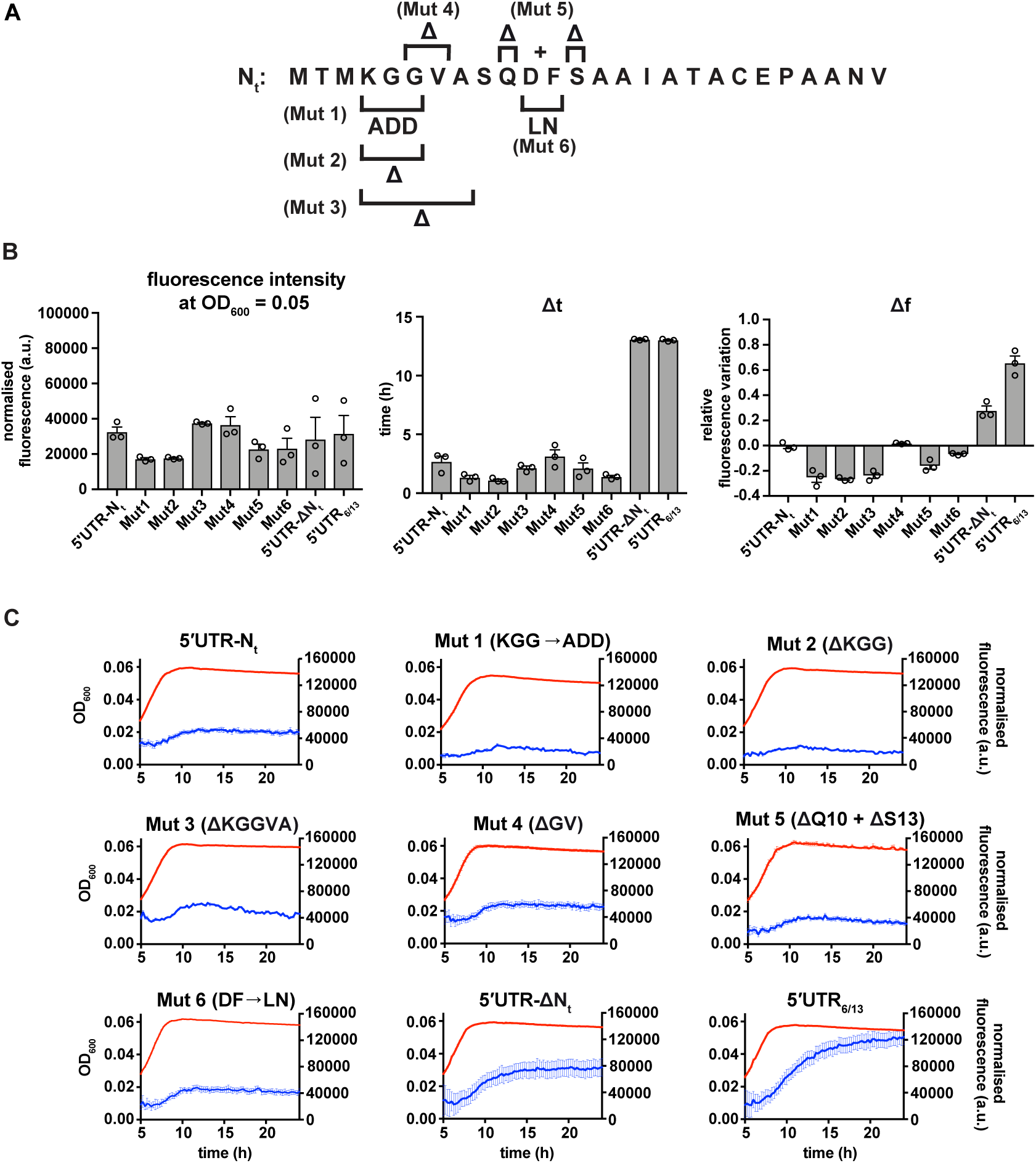
Additional mutations in the N_t_ amino acid sequence. (**A**) Schematic illustration mutations Mut 1-6. (**B**) Values of fluorescence intensity at OD_600_ = 0.05, Δt and Δf calculated for the mutant reporter strains illustrated in (A). (**C**) Growth curves and fluorescence kinetic profiles of reporter strains shown in (A), with profiles of the 5’UTR-N_t_ and 5’UTR_6/13_ strains for comparison. Averages of three independent replicates are shown with error bars representing standard errors.

**Supplementary Fig. 9.**
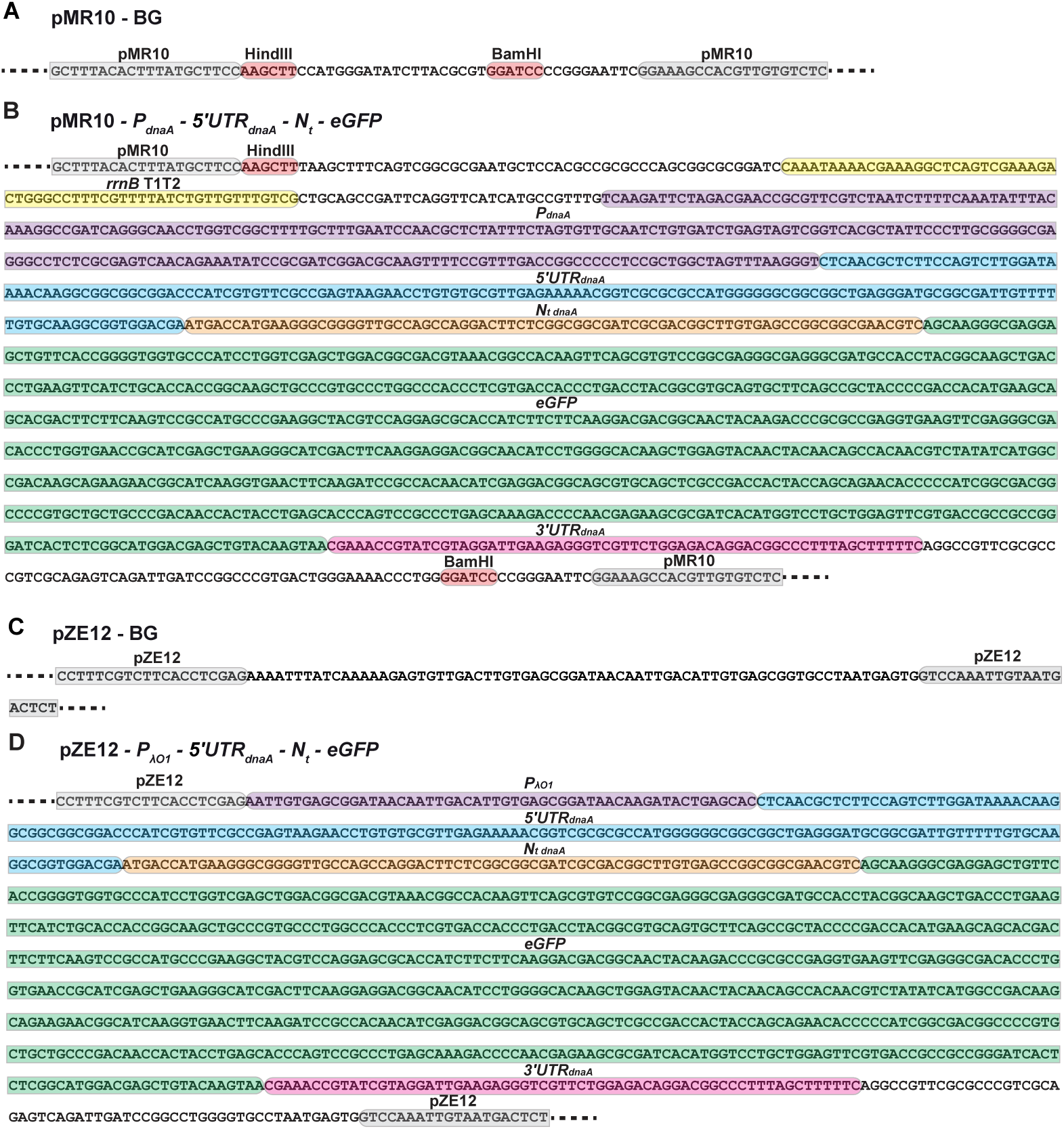
pMR10 and pZE12 reporter plasmid construction. (**A**) Sequence of the multicloning site region in the new pMR10-BG construct derived from plasmid pMR10. Grey: flanking sequences of the parental pMR10 plasmid. (**B**) The pMR10-*P_dnaA_*-5’UTR*_dnaA_*-N_t_-eGFP plasmid was derived from the construct in (A) upon digestion of pMR10-BG with HindIII and BamHI (restriction sites in red). The insert comprised, the *rrnB1*-T1T2 terminator (yellow), 245 bp upstream of the *dnaA* transcription start site (*P_dnaA_* - purple), the 5’UTR*_dnaA_* (blue), the first 78 bp of *dnaA* open reading frame (orange), the eGFP gene (green) and 63 bp downstream of the *dnaA* stop codon (3’UTR*_dnaA_* - pink). (**C**) Plasmid pZE12-BG was derived from the pZE12-luc plasmid by deleting the luciferase gene and part of the *P_λO1_* promoter. In grey, the flanking sequences of the original pZE12-luc plasmid. (**D**) The pZE12-*P*_λO1_-5’UTR*_dnaA_*-N_t_-eGFP reporter plasmid, was obtained from the pZE12-luc plasmid by exchanging the luciferase gene downstream of the Lambda O1 promoter (*P_λO1_* – purple) with an insert containing the 5’UTR*_dnaA_* (blue), the first 78 nt of *dnaA* open reading frame (orange), the eGFP gene (green) and the 3’UTR*_dnaA_* (pink).

**Supplementary Table 1.**
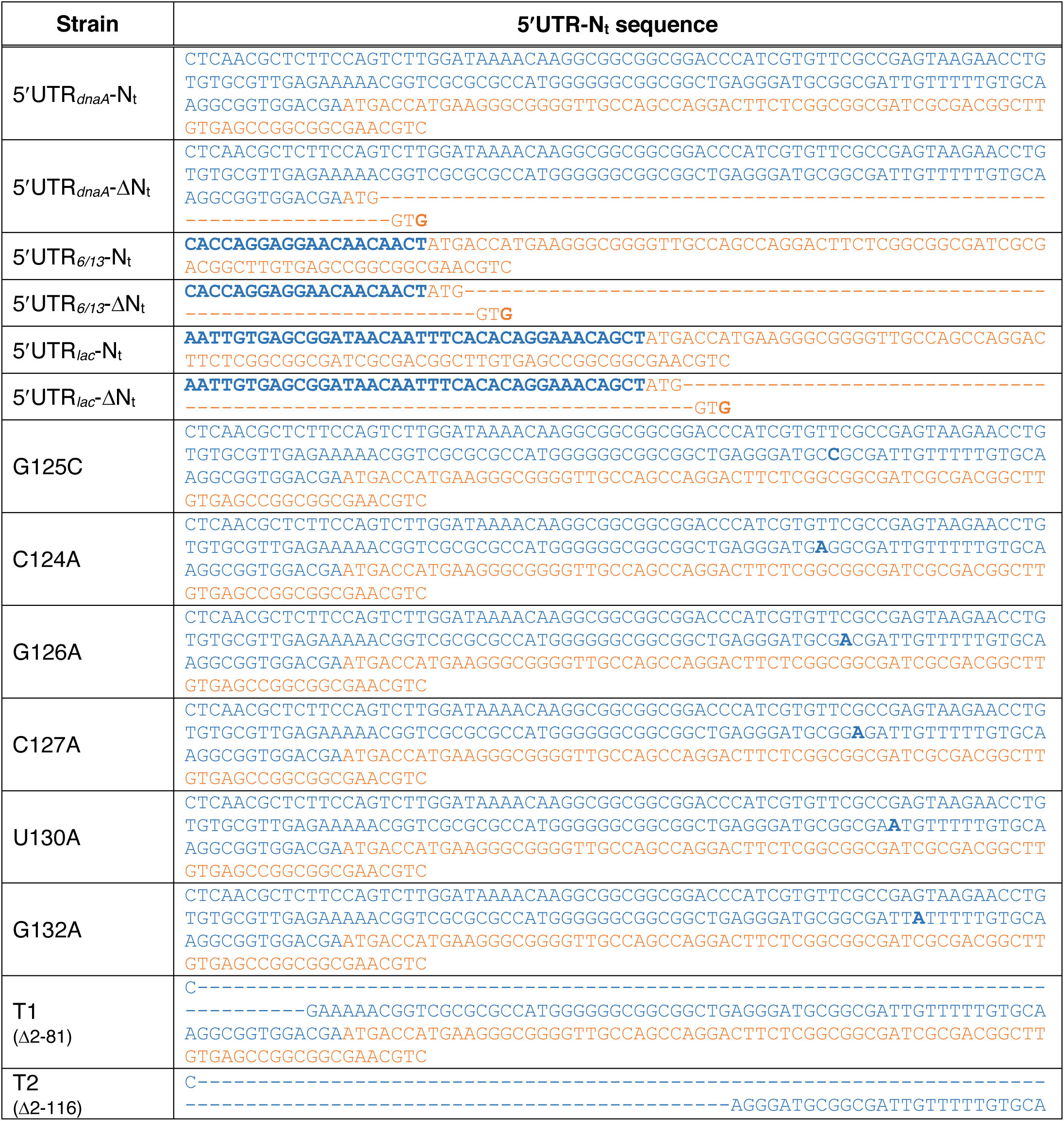
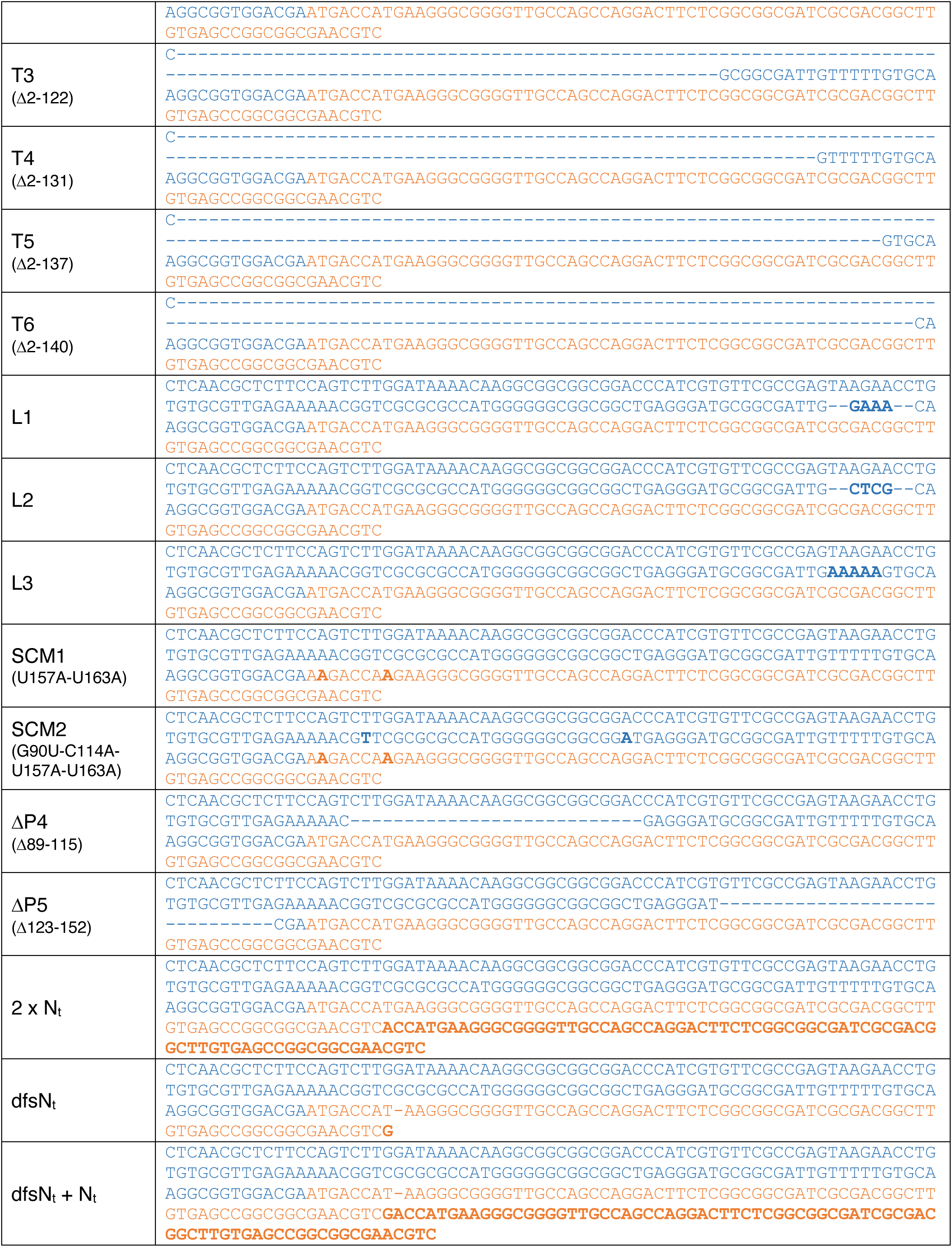
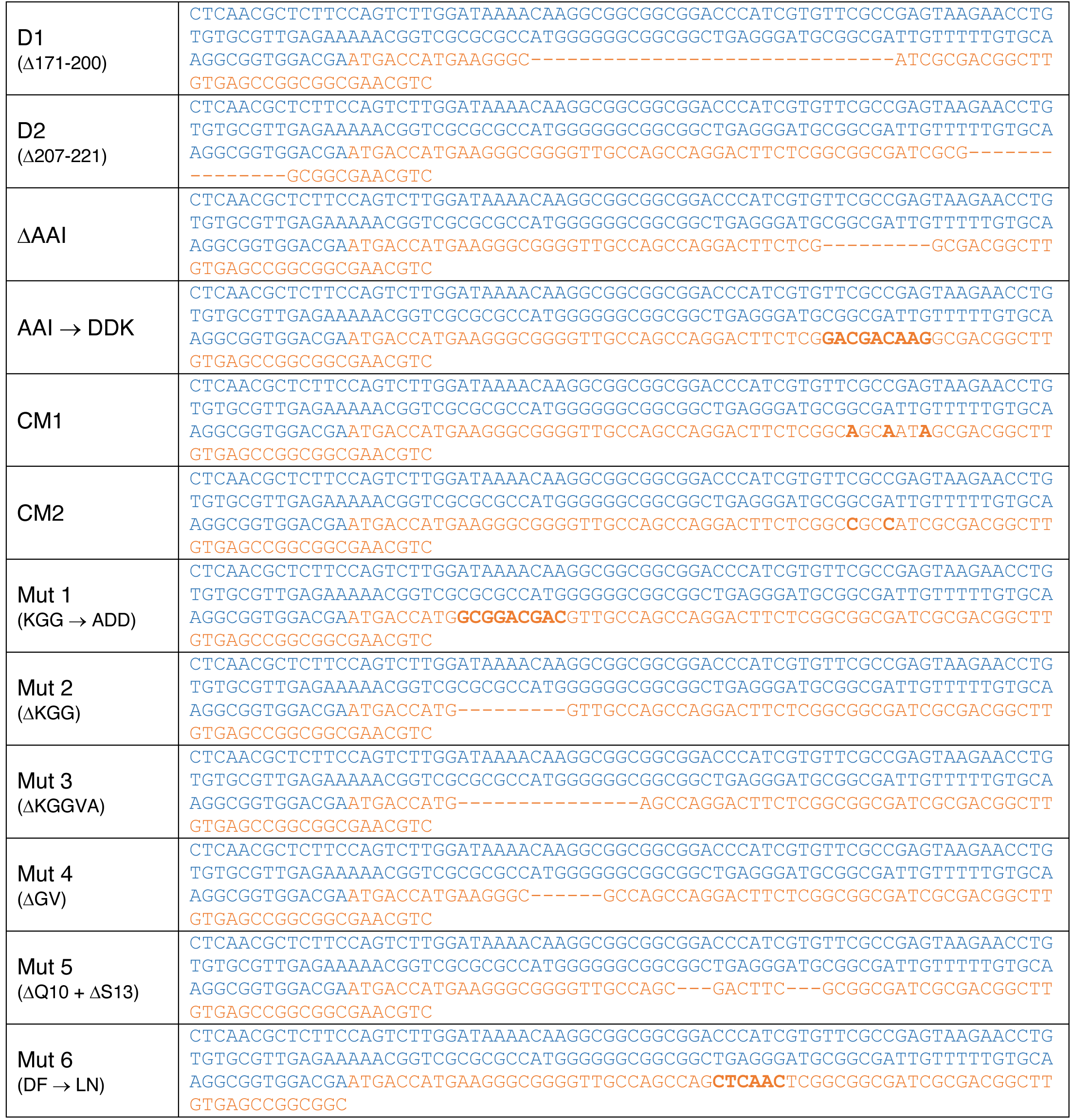
Sequence of the 5*′*UTR and N_t_ modules in the reporter plasmid constructs. The reporter plasmids were generated by site-directed mutagenesis, using the 5*′*UTR*_dnaA_*-N_t_ construct as a PCR template (Supplementary Fig. 9B and 9D). The sequences of the 5*′*UTR and N_t_ regions are shown in blue and orange respectively. Nucleotide substitutions and insertions are indicated in bold. Nucleotide deletions are indicated with the symbol “–”.

**Supplementary Table 2.** List of bacterial strains used in this study. The sequences of the 5*′*UTR and N_t_ regions are shown in the Supplementary Table 1.

| Bacterial strain / plasmid | Genetic background | Source |
| --- | --- | --- |
| pMR10-BG | *Caulobacter crescentus* NA1000 | This study |
| pMR10-5'UTR<sub>dnaA</sub>-N<sub>t</sub> | *Caulobacter crescentus* NA1000 | This study |
| pMR10-5'UTR<sub>dnaA</sub>-ΔN<sub>t</sub> | *Caulobacter crescentus* NA1000 | This study |
| pMR10-5'UTR<sub>6/13</sub>-N<sub>t</sub> | *Caulobacter crescentus* NA1000 | This study |
| pMR10-5'UTR<sub>6/13</sub>-ΔN<sub>t</sub> | *Caulobacter crescentus* NA1000 | This study |
| pMR10-5'UTR<sub>lac</sub>-N<sub>t</sub> | *Caulobacter crescentus* NA1000 | This study |
| pMR10-5'UTR<sub>lac</sub>-ΔN<sub>t</sub> | *Caulobacter crescentus* NA1000 | This study |
| pMR10-G125C | *Caulobacter crescentus* NA1000 | This study |
| pMR10-C124A | *Caulobacter crescentus* NA1000 | This study |
| pMR10-G126A | *Caulobacter crescentus* NA1000 | This study |
| pMR10-C127A | *Caulobacter crescentus* NA1000 | This study |
| pMR10-U130A | *Caulobacter crescentus* NA1000 | This study |
| pMR10-G132A | *Caulobacter crescentus* NA1000 | This study |
| pMR10-T1 | *Caulobacter crescentus* NA1000 | This study |
| pMR10-T2 | *Caulobacter crescentus* NA1000 | This study |
| pMR10-T3 | *Caulobacter crescentus* NA1000 | This study |
| pMR10-T4 | *Caulobacter crescentus* NA1000 | This study |
| pMR10-T5 | *Caulobacter crescentus* NA1000 | This study |
| pMR10-T6 | *Caulobacter crescentus* NA1000 | This study |
| pMR10-L1 | *Caulobacter crescentus* NA1000 | This study |
| pMR10-L2 | *Caulobacter crescentus* NA1000 | This study |
| pMR10-L3 | *Caulobacter crescentus* NA1000 | This study |
| pMR10-SCM1 | *Caulobacter crescentus* NA1000 | This study |
| pMR10-SCM2 | *Caulobacter crescentus* NA1000 | This study |
| pMR10-ΔP4 | *Caulobacter crescentus* NA1000 | This study |
| pMR10-ΔP5 | *Caulobacter crescentus* NA1000 | This study |
| pMR10-2xN<sub>t</sub> | *Caulobacter crescentus* NA1000 | This study |
| pMR10-dfsN<sub>t</sub> | *Caulobacter crescentus* NA1000 | This study |
| pMR10-dfsN<sub>t</sub> + N<sub>t</sub> | *Caulobacter crescentus* NA1000 | This study |
| pMR10-D1 | *Caulobacter crescentus* NA1000 | This study |
| pMR10-D2 | *Caulobacter crescentus* NA1000 | This study |
| pMR10-ΔAAI | *Caulobacter crescentus* NA1000 | This study |
| pMR10-AAI-DDK | *Caulobacter crescentus* NA1000 | This study |
| pMR10-CM1 | *Caulobacter crescentus* NA1000 | This study |
| pMR10-CM2 | *Caulobacter crescentus* NA1000 | This study |
| pMR10-Mut 1 | *Caulobacter crescentus* NA1000 | This study |
| pMR10-Mut 2 | *Caulobacter crescentus* NA1000 | This study |
| pMR10-Mut 3 | *Caulobacter crescentus* NA1000 | This study |
| pMR10-Mut 4 | *Caulobacter crescentus* NA1000 | This study |
| pMR10-Mut 5 | *Caulobacter crescentus* NA1000 | This study |
| pMR10-Mut 6 | *Caulobacter crescentus* NA1000 | This study |
| pZE12-BG | *Escherichia coli* MG1655 | This study |
| pZE12-5'UTR<sub>dnaA</sub>-N<sub>t</sub> | *Escherichia coli* MG1655 | This study |
| pZE12-5'UTR<sub>dnaA</sub>-ΔN<sub>t</sub> | *Escherichia coli* MG1655 | This study |
| pZE12-5'UTR<sub>6/13</sub>-N<sub>t</sub> | *Escherichia coli* MG1655 | This study |
| pZE12-5'UTR<sub>6/13</sub>-ΔN<sub>t</sub> | *Escherichia coli* MG1655 | This study |

## Notes

### Competing Interest Statement

The authors have declared no competing interest.

